# Extracellular contractile injection systems for receptor-mediated protein delivery into plant cells

**DOI:** 10.1101/2025.08.22.671880

**Authors:** Mark G. Legendre, Carlos A. Heredia, Clair Colee, Kimberley T. Muchenje, Yunqing Wang, Virginia Pistilli, Gozde S. Demirer

## Abstract

Protein delivery into plant cells remains a major barrier in plant biotechnology. Extracellular contractile injection systems (eCIS) are phage tail-like nanomachines evolved by bacteria to engage eukaryotic host cells and deliver protein effectors. The *Photorhabdus* virulence cassette (PVC) is a well-characterized eCIS that naturally targets insect hosts. Here, we retargeted PVCs to plants by engineering their tail fibers to recognize the plant receptor FLAGELLIN SENSITIVE2 (FLS2). We designed and characterized a library of FLS2-targeting PVC variants and evaluated their ability to deliver native and non-native protein cargoes, including a fluorescent protein, Cre recombinase, and R2 retroelement genome editor. Engineered PVCs delivered Cre into wild-type *Arabidopsis thaliana* protoplasts and *Nicotiana benthamiana* leaf cells with efficiencies of up to 15%. Delivery was FLS2-dependent and scaled with receptor abundance, reaching more than 75% of leaf cells in FLS2-overexpressing lines. We further demonstrated PVC-mediated delivery of R2, resulting in targeted insertion of an mCherry reporter into the plant genome. This work establishes PVCs as receptor-mediated protein-delivery vehicles for plant cells and provides a foundation for targeted, DNA-free genome engineering in plant biotechnology.

## Introduction

Advances in plant biotechnology position genetic engineering as a sustainable route to improved crops^1^. Direct protein delivery is particularly attractive because protein cargoes act immediately after cell entry, avoid transcriptional and translational delays, and are transient by design. For genome editing, delivery of preformed editors can eliminate nuclease-encoding DNA and restrict the duration of editor activity, reducing risks associated with unwanted transgene integration and prolonged editor expression^2^. Yet, proteins are challenging cargoes: they are often large and structurally complex, must remain folded and active, are vulnerable to proteolytic degradation, and generally cannot be packaged by the electrostatic complexation strategies used for nucleic acids.

Although nanoparticles have advanced nucleic acid delivery to plants^3–12^, protein delivery tools remain underdeveloped. Current methods include particle bombardment, in which protein-coated microparticles are physically propelled into the explant tissue but can cause substantial mechanical damage and lack cellular targeting^13^; cell-penetrating peptides, which form complexes with protein cargoes and promote membrane translocation but are often low-efficiency^14–21;^ and a recently reported study of vehicle-free protein internalization, which can deliver genome-editing proteins into walled cells and whole-plant tissues in some contexts^22^. These approaches demonstrate the promise and highlight the need for protein delivery platforms that combine efficient intracellular delivery, compatibility with intact plant tissues, and programmable cellular targeting.

Protein nanoparticles offer an alternative strategy for protein delivery because they can be genetically encoded, self-assemble into defined architectures, and, in some cases, package and deliver protein cargoes through evolved mechanisms^23,24^. In nature, many bacteria use specialized protein nanomachines to translocate effector proteins into eukaryotic host cells. Contractile injection systems are a prominent example: phage tail-like nanosyringes composed of a rigid inner tube tipped by a spike, surrounded by a contractile sheath, and anchored to a baseplate^25–28^. Following target-cell recognition, sheath contraction propels the inner tube across the target membrane, delivering the loaded protein payloads directly into the host cytosol^29–32^.

Contractile injection systems can remain anchored to bacterial membranes, where delivery requires direct cell-to-cell contact^33^, or can be released as extracellular complexes after lysis of the producing cell. These extracellular contractile injection systems (eCIS) function independently of their bacterial hosts and interact with a diverse set of target cells to deliver protein cargoes^34–37^. Structural and biochemical studies have revealed mechanisms of selective cargo loading^38–40^, while engineering studies have shown that tail-fiber modification can alter host specificity^41–43^. More recently, engineered eCIS have been used to deliver functional protein cargoes, including genome-editing machinery, into mammalian cells^38,44^. Intraluminal engineering has further expanded eCIS cargo capacity to RNA, enabling co-delivery of Cas nucleases and guide RNAs that assemble into functional ribonucleoproteins for *in vivo* genome editing in mouse tissues^45^.

*Photorhabdus* virulence cassette (PVC) is among the best-characterized eCIS subfamilies^40,46^. PVCs naturally deliver toxin effectors into insect host cells, and recent studies have retargeted them to deliver heterologous proteins into mammalian cells^44,47^. Yet, PVCs have not been adapted for use in plants. Receptor-mediated protein delivery could address a central bottleneck in plant biotechnology by enabling targeted delivery of proteins, including genome-engineering cargoes, while avoiding delivery of nuclease-encoding DNA or other transgene-encoding constructs.

Here, we harnessed PVC extracellular contractile injection systems for receptor-mediated protein delivery into plant cells by reprogramming their tail fibers to recognize the plant receptor FLAGELLIN SENSITIVE2 (FLS2). We designed and characterized a library of FLS2-targeting PVC variants, established receptor-dependent delivery using FLS2 knockout and overexpression lines, and showed that delivery scales with receptor abundance, reaching more than 75% of *Arabidopsis* leaf cells in overexpression lines. We demonstrated the delivery of multiple functional cargoes, including a fluorescent protein, a Cre recombinase, and the R2 retroelement genome editor, enabling targeted gene insertion at the plant ribosomal DNA locus. Together, these results establish retargeted PVCs as programmable protein-delivery vehicles for plant cells and provide a foundation for DNA-free delivery of genome-engineering proteins in plant biotechnology.

## Results

### The PVC eCIS can be retargeted to the plant cell membrane

The PVC gene cluster consists of 16 structural genes (*pvc1-16*) that are responsible for the expression and self-assembly of the functional nanoparticle^48^. Immediately downstream of these structural genes are two payload genes, *pnf* and *pdp1*, which encode native toxin effector proteins. In addition to these two functional cargo genes, four accessory genes are necessary for cargo loading. Combined, these structural and payload genes constitute the *PVCpnf* gene locus (**Fig. 1A**).

**Figure 1.**
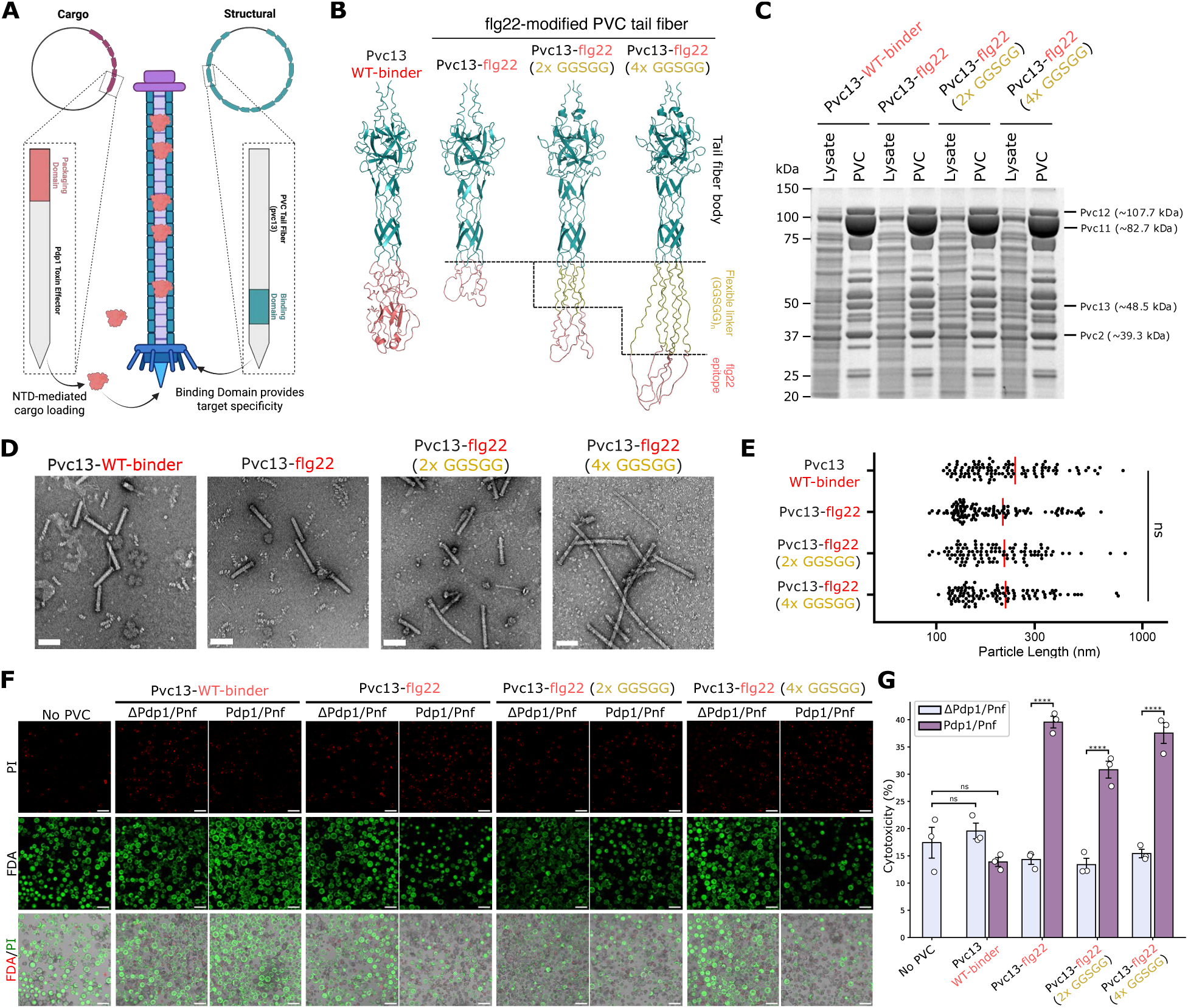
Rational design of PVCs targeting the plant receptor FLS2. **(A)** Constructs to express PVCs in *E. coli*: a structural plasmid containing all 16 structural genes (teal) necessary for particle formation, and a cargo plasmid with four accessory genes and a single cargo-coding sequence (salmon). **(B)** AlphaFold-predicted structures of a trimeric Pvc13 tail fiber library designed to target FLS2. Only the terminal region of the tail fiber homotrimer is visualized, containing part of the tail fiber body (teal), an engineered flexible linker (yellow), and the flg22 epitope (salmon). **(C)** SDS-PAGE of PVC libraries shows structural proteins within the PVC particle, including baseplate proteins Pvc11 and Pvc12, sheath protein Pvc2, and engineered tail fibers Pvc13. **(D)** Negative-stain TEM images of PVC library in their non-ejected states. Scale bars, 100 nm. **(E)** Particle-length profiles of all PVC variants loaded with toxin cargoes from TEM images. **(F)** Representative confocal microscopy images of protoplasts 24 h after challenge with PVCs containing either the native insect-targeting tail fiber (Pvc13-WT-binder) or FLS2-targeting variants. PVC treatments include toxin-loaded particles (Pdp1/Pnf) or empty particles (ΔPdp1/Pnf). Scale bars, 100 µm. **(G)** Quantification of cytotoxicity for all PVC variants and a WI buffer control (No PVC). Cytotoxicity is expressed as the number of PI-stained cells relative to the total number of cells (PI + FDA). Data are mean ± SEM with n = 3 biological replicates; one-way ANOVA with Tukey post hoc test. ****p < 0.0001. ns, not significant.

*E. coli* has previously been engineered to recombinantly express the *PVCpnf* locus from *P. asymbiotica*^44^. Here, we used a modular two-plasmid expression system in which the *PVCpnf* locus was divided into structural and cargo vectors, termed pStructural and pCargo, respectively (**Fig. 1A**). PVC generation and purification were achieved using previously described methods^48^, in which co-transformation of pStructural and pCargo into *E. coli* produced defined PVC variants with specified tail-fiber and cargo configurations.

The PVC tail-fiber binding domain has been identified and engineered to redirect PVC tropism toward mammalian hosts^44^. We tested whether an analogous strategy could generate PVCs that recognize plant cell-surface receptors and deliver protein cargoes into plant cells. We used AlphaFold2 to predict the structure of the distal tip of the tail fiber encoded by *pvc13* (**Fig. 1B**). The tail fiber was predicted to assemble as a Pvc13 homotrimer, with a helical tail-fiber body and a terminal globular region corresponding to the previously identified receptor-binding domain^44^.

For retargeting, we selected FLS2 because it is a highly expressed, cell-surface-localized, and extensively characterized plant receptor with a structurally defined peptide ligand, making it a tractable proof-of-concept target for developing receptor-mediated PVC delivery in plants. FLS2 is a microbe-associated molecular pattern receptor that recognizes flg22, a conserved 22-amino-acid epitope from bacterial flagellin^49,50^. We engineered the Pvc13 binding domain to display flg22, with the goal of redirecting PVC binding toward FLS2 at the plant plasma membrane.

We designed a PVC library displaying flg22 at different distances from the tail-fiber body using (GGSGG)_n=0,2,4_ flexible linkers at each peptide terminus to reduce potential steric constraints on receptor binding. These particles are named as Pvc13-flg22, Pvc13-flg22 (2xGGSGG), and Pvc13-flg22 (4xGGSGG), respectively. AlphaFold predictions suggested that all variants preserved the expected helical tail-fiber body while maintaining the predicted disorder of the displayed flg22 epitope (**Fig. 1B**). ChimeraX alignment of each FLS2-targeting PVC variant to the FLS2 binding region further suggested that the epitope could adopt the expected receptor-binding conformation independent of linker length and despite fusion to the larger tail-fiber scaffold (**Fig. S1**).

After expression and purification, denaturing gel electrophoresis verified incorporation of the expected PVC structural proteins into each member of the PVC library (**Fig. 1C**). Negative-stain transmission electron microscopy (TEM) confirmed self-assembly of nanoparticles loaded with the native toxin cargoes (**Fig. 1D**). All engineered variants retained the expected PVC morphology and showed similar particle-length distributions, indicating that tail-fiber modification did not disrupt particle assembly (**Fig. 1E**). Self-assembly was maintained across several physiological buffers; however, buffer-dependent destabilization and spontaneous sheath contraction constrained the formulations in which these nanoparticles could be applied (**Fig. S2**).

Circular dichroism (CD) analysis further showed that tail-fiber engineering did not detectably alter PVC structure or stability. Far-UV and near-UV thermal melts showed highly similar secondary structure and higher-order nanoparticle stability across all PVC library members, with melting temperatures of 55-60°C and 50-52°C, respectively **(Fig. S3**). Together, these biochemical, TEM, and CD analyses confirm the production and isolation of structurally and functionally intact recombinant PVC nanoparticles for plant targeting.

To test whether FLS2-targeting PVCs could deliver proteins into plant cells, we loaded them with the native insect toxin effectors Pnf and Pdp1 and applied the particles to *Arabidopsis thaliana* protoplasts. All toxin-loaded FLS2-targeting PVCs caused significantly greater cell mortality than the native insect-targeting Pvc13-WT-binder, as measured by PI/FDA live-dead staining (**Fig. 1F, G**). In contrast, empty FLS2-targeting PVCs (ΔPdp1/Pnf) did not induce cell death above background (**Fig. 1F, G**), indicating that cytotoxicity was cargo-dependent rather than caused by nonspecific PVC toxicity. These results provide initial evidence that FLS2-targeting PVCs can deliver protein cargoes into plant cells.

### PVC-mediated delivery of non-native protein cargoes to plant cells

Although the molecular mechanism of PVC cargo loading remains incompletely understood, N-terminal sequences in the native Pnf and Pdp1 effectors are known to mediate cargo loading into the PVC sheath lumen^38,39^. We developed a cargo-engineering strategy in which the Pdp1 N-terminal packaging domain (NTD) is fused to the N-terminus of a non-native cargo protein, and a HiBiT tag is fused to the C-terminus for chemiluminescent detection (**Fig. 2A**). This design enables loading and detection of engineered cargoes and can accommodate additional tags, such as for affinity purification, protein stability, and subcellular localization.

**Figure 2.**
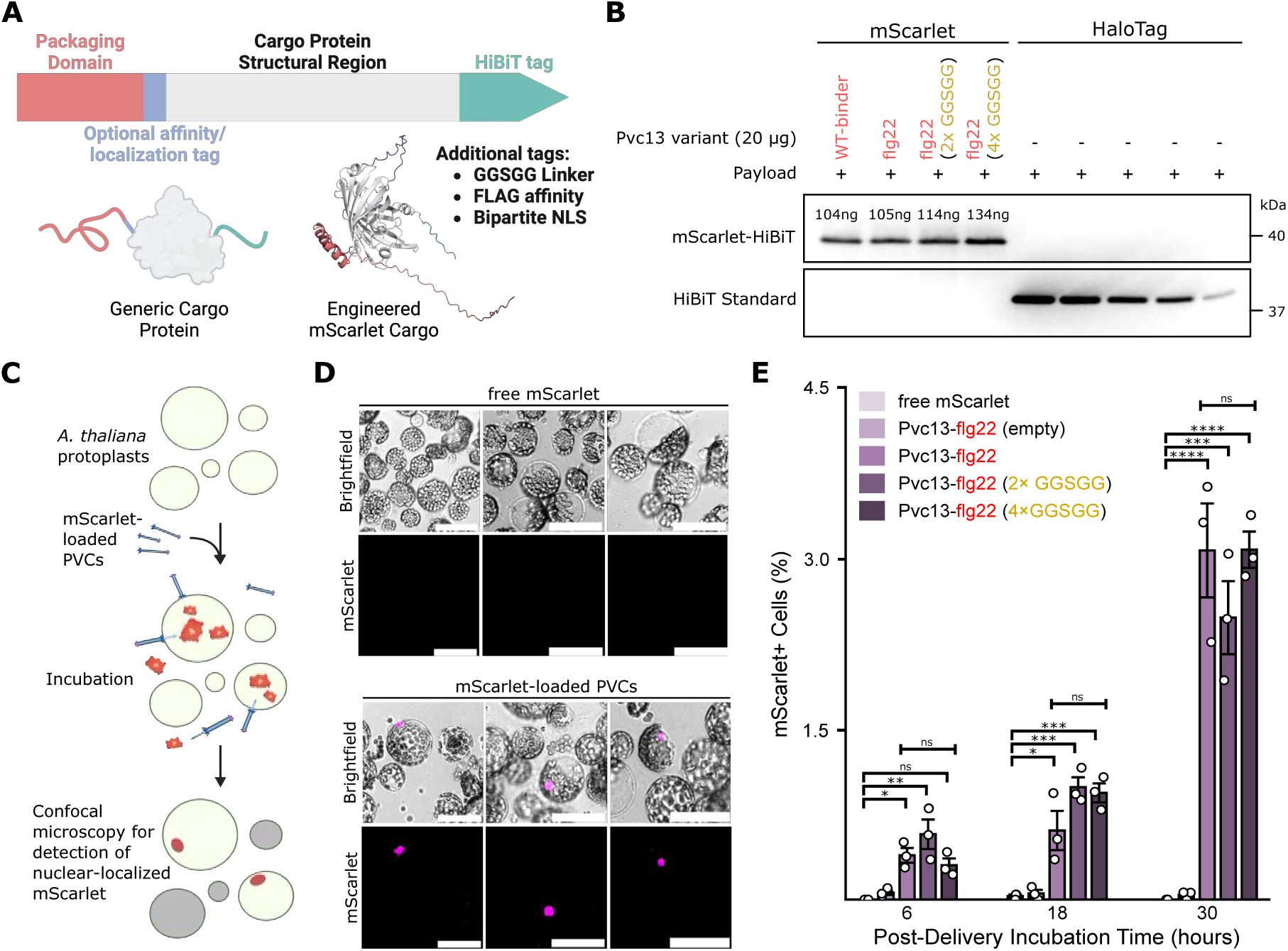
PVC-mediated delivery of a fluorescent protein to protoplasts. (**A)** Engineering strategy for loading non-native cargoes involves tagging the cargo protein N-terminus with the *Pdp1* packaging domain and its C-terminus with a HiBiT tag. The design strategy accommodates additional tagging and is applied to an NLS-fused mScarlet fluorescent protein. **(B)** Denaturing Western blot analysis targeting the HiBiT tag on the mScarlet cargo loaded in all PVC variants. mScarlet loading quantification was performed against a HiBiT standard. **(C)** Schematic of the PVC-mediated fluorescent protein delivery workflow in *A. thaliana* protoplasts. **(D)** Representative confocal microscopy images of *A. thaliana* protoplasts challenged with mScarlet-loaded Pvc13-flg22 or a recombinant NLS-fused mScarlet control at an equivalent cargo amount. Scale bars, 50 µm. **(E)** Quantification of %mScarlet+ cells for all PVC variants and controls across 6 to 30 h incubation. Data are mean ± SEM with n = 3 biological replicates; one-way ANOVA with Tukey post hoc test. *p < 0.05, **p < 0.01, ***p < 0.001, ****p < 0.0001. ns, not significant.

Using this strategy, we loaded fluorescent protein mScarlet into PVCs, generating a 39.8-kDa engineered cargo protein. In addition to NTD and HiBiT tags, mScarlet contained an N-terminal bipartite nuclear-localization sequence (NLS) to concentrate signal in the nucleus and facilitate microscopy-based detection (**Fig. 2A**). A denaturing Western blot against the HiBiT tag confirmed mScarlet loading into PVCs, with an estimated loading of 4.5 ng mScarlet per 1 µg of Pvc13-flg22 (**Fig. S4A**). mScarlet loading was similar across linker variants, with comparable cargo levels at matched PVC concentrations (**Fig. 2B, Fig. S4B**). TEM analysis showed that non-native cargo loading did not detectably disrupt nanoparticle structural integrity, with only minor differences in particle-length distributions across structural variants carrying different cargoes (**Fig. S5**).

We next tested whether mScarlet-loaded FLS2-targeting PVCs could deliver into plant cells. *A. thaliana* mesophyll protoplasts were treated with mScarlet-loaded Pvc13-flg22 or recombinant NLS-fused mScarlet alone at an equivalent cargo concentration (4.5 ng mL⁻¹) (**Fig. 2C, Fig. S4C**). Nuclear mScarlet signal was detected only in PVC-mediated delivery and not in treatment with recombinant mScarlet alone (**Fig. 2D**). Across all FLS2-targeting PVC variants, delivery increased from 6 to 30 h, with mScarlet-loaded PVCs significantly outperforming both free mScarlet and empty PVC controls at all time points (**Fig. 2E, Fig. S4**). Although delivery increased with longer incubation, all FLS2-targeting linker variants performed similarly (**Fig. 2E**). These data show PVC-mediated delivery of a non-native protein cargo into plant cells.

### PVCs can deliver Cre recombinase to plant protoplasts

After confirming mScarlet delivery, we next tested Cre recombinase. Cre has a lower molecular weight than the native Pnf cargo, making it a suitable initial genome-engineering cargo. We first developed an *in planta* reporter assay for detecting intracellular Cre activity, using a Cre flip-excision (FLEX) reporter^51^ containing an inverted N7 NLS-tagged YFP coding sequence flanked by two orthogonal pairs of *loxP* sites (**Fig. S6A**). Cre activity irreversibly inverts the reporter to produce a nuclear YFP signal. We generated two reporter versions: a standard FLEX reporter and a geminiviral replicon-amplified FLEX-GV reporter designed to increase reporter copy number, sensitivity, and signal output (**Fig. S6B**). Both reporters showed minimal leakiness in negative controls and robust activation when co-delivered with a constitutive Cre expression plasmid in *A. thaliana* protoplasts and *N. benthamiana* leaves (**Fig. S6C, D**). These results establish two Cre-responsive reporters to detect the intracellular activity of exogenously delivered Cre *in planta*.

Using the cargo-engineering strategy described above, we fused the packaging domain to the N-terminus of bipartite NLS-tagged Cre and added a C-terminal HiBiT tag for chemiluminescent cargo detection (**Fig. 3A**). Denaturing Western blot analysis confirmed Cre loading into PVCs at ∼10 ng Cre per 1 µg PVC, with comparable loading across all PVC variants (**Fig. 3B and Fig. S7**).

**Figure 3.**
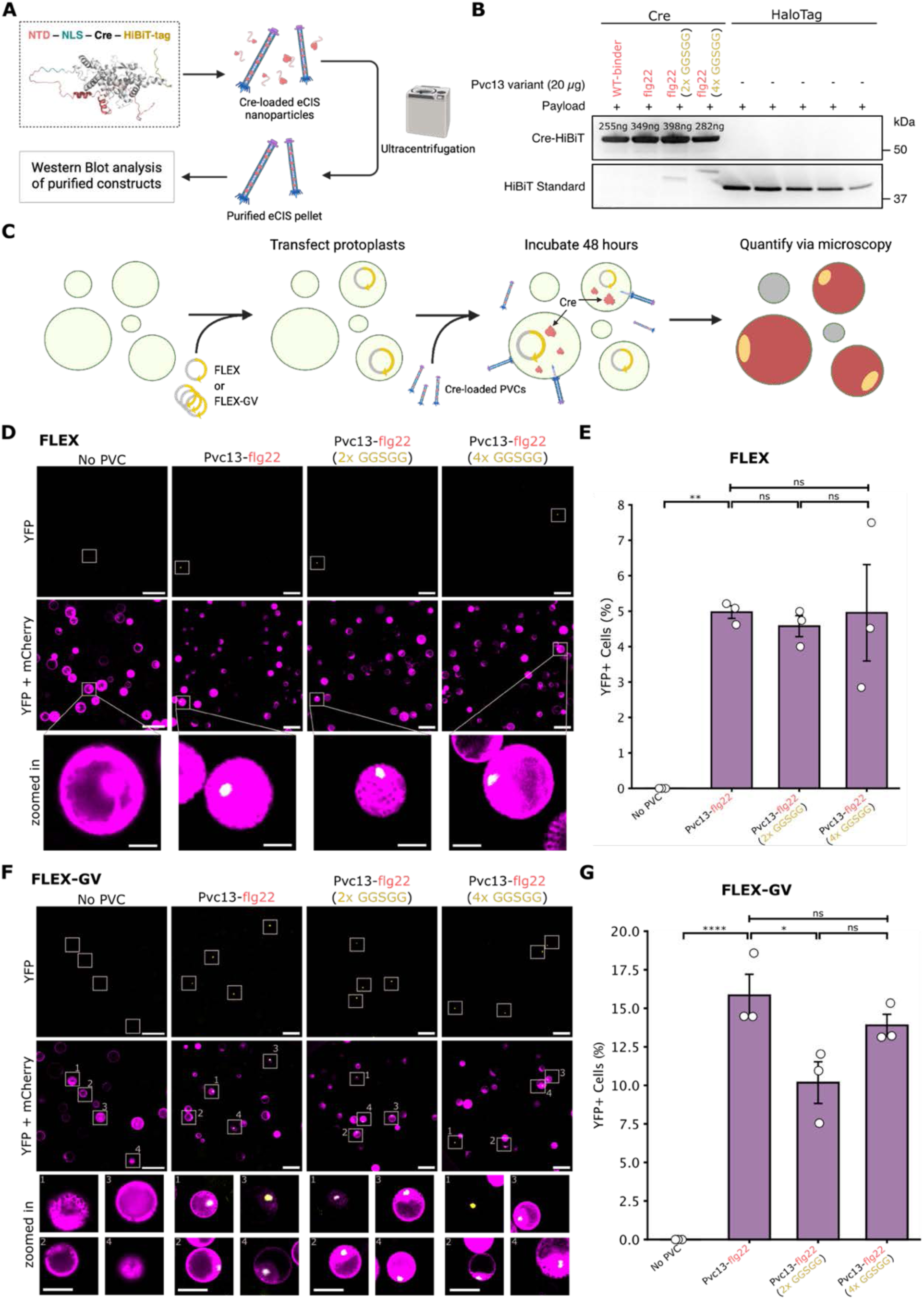
Reprogrammed PVCs deliver Cre protein cargoes to plant protoplasts. **(A)** Cre recombinase with the N-terminal packaging domain and a bipartite NLS, and a C-terminal HiBiT tag. **(B)** Denaturing Western blot analysis of the cargo HiBiT tag visualizes the presence of non-native Cre in all PVC variants. Quantification of Cre recombinase loading was performed against a HiBiT standard. **(C)** Schematic of the PVC delivery workflow in *Arabidopsis thaliana* protoplasts, where successful delivery events are registered as nuclear YFP fluorescence. **(D)** Representative confocal fluorescence microscopy images of *A. thaliana* protoplasts transfected with the FLEX reporter and treated with the Cre-loaded FLS2-targeting PVC library. Scale bar, 100 µm. Inset, 40 µm. **(E)** Quantification of %YFP+ cells from confocal images in panel D using the FLEX reporter. **(F)** Representative confocal fluorescence microscopy images of *A. thaliana* protoplasts transfected with the FLEX-GV reporter and treated with the Cre-loaded FLS2-targeting PVC library. Scale bar, 100 µm. Inset, 40 µm. **(G)** Quantification of %YFP+ cells from confocal images in panel F using the FLEX-GV reporter. Data are mean ± SEM with n = 3 biological replicates; one-way ANOVA with Tukey post hoc test. *p < 0.05, **p < 0.01, and ****p < 0.0001. ns, not significant.

We delivered Cre-loaded Pvc13-flg22 to FLEX reporter-expressing *A. thaliana* protoplasts and optimized assay conditions. YFP signal peaked as early as 24 h (**Fig. S7A, B**), and the % YFP+ cells increased with PVC concentration up to 500 ng µL⁻¹ (**Fig. S7C, D**). We used these optimal delivery conditions to screen the Cre-loaded FLS2-binding PVC library in *A. thaliana* protoplasts (**Fig. 3C**). With the standard FLEX reporter, all FLS2-targeting variants produced similar reporter activity, with ∼5% YFP+ cells (**Fig. 3D, E**). Delivery required both FLS2-targeting and Cre, as neither WT-binder particles nor empty Pvc13-flg22 particles produced significant reporter activity (**Fig. S7E, F**). The higher-sensitivity FLEX-GV reporter increased detectable delivery to 16%, 10%, and 15% for Pvc13-flg22, Pvc13-flg22 (2×GGSGG), and Pvc13-flg22 (4×GGSGG), respectively (**Fig. 3F, G**).

We benchmarked Pvc13-flg22 PVCs against PEG-mediated protein transfection, a common protoplast protein delivery method. Under our assay conditions, PVC-mediated Cre delivery was statistically comparable to PEG-mediated Cre delivery using the FLEX-GV reporter, supporting PVCs as a competitive protein-delivery platform in protoplasts (**Supplementary Fig. 6G, H**).

### PVCs can deliver Cre recombinase to walled leaf cells

We next tested Cre-loaded FLS2-targeting PVCs in *N. benthamiana* leaves expressing FLEX reporters (**Fig. 4A**). High-resolution TEM revealed PVC nanoparticles in plant tissue adjacent to and within the cell wall, supporting the physical association of PVCs with walled plant cells prior to delivery (**Fig. S8**). Time-course and dose-response analyses showed optimal performance at PVC concentrations above 200 ng µL⁻¹ and after 3 days of incubation across linker variants (**Fig. S9**). After 72 h of PVC incubation, Cre-driven YFP expression was low when detected with the low-copy FLEX reporter, likely due to limited reporter sensitivity (**Fig. 4B, C**). In contrast, the FLEX-GV reporter detected YFP expression in 13%, 16%, and 10% of leaf cells treated with Pvc13-flg22, Pvc13-flg22 (2×GGSGG), and Pvc13-flg22 (4×GGSGG), respectively (**Fig. 4D, E**), comparable to protoplast delivery efficiencies detected with the FLEX-GV reporter (**Fig. 3F, G**).

**Figure 4.**
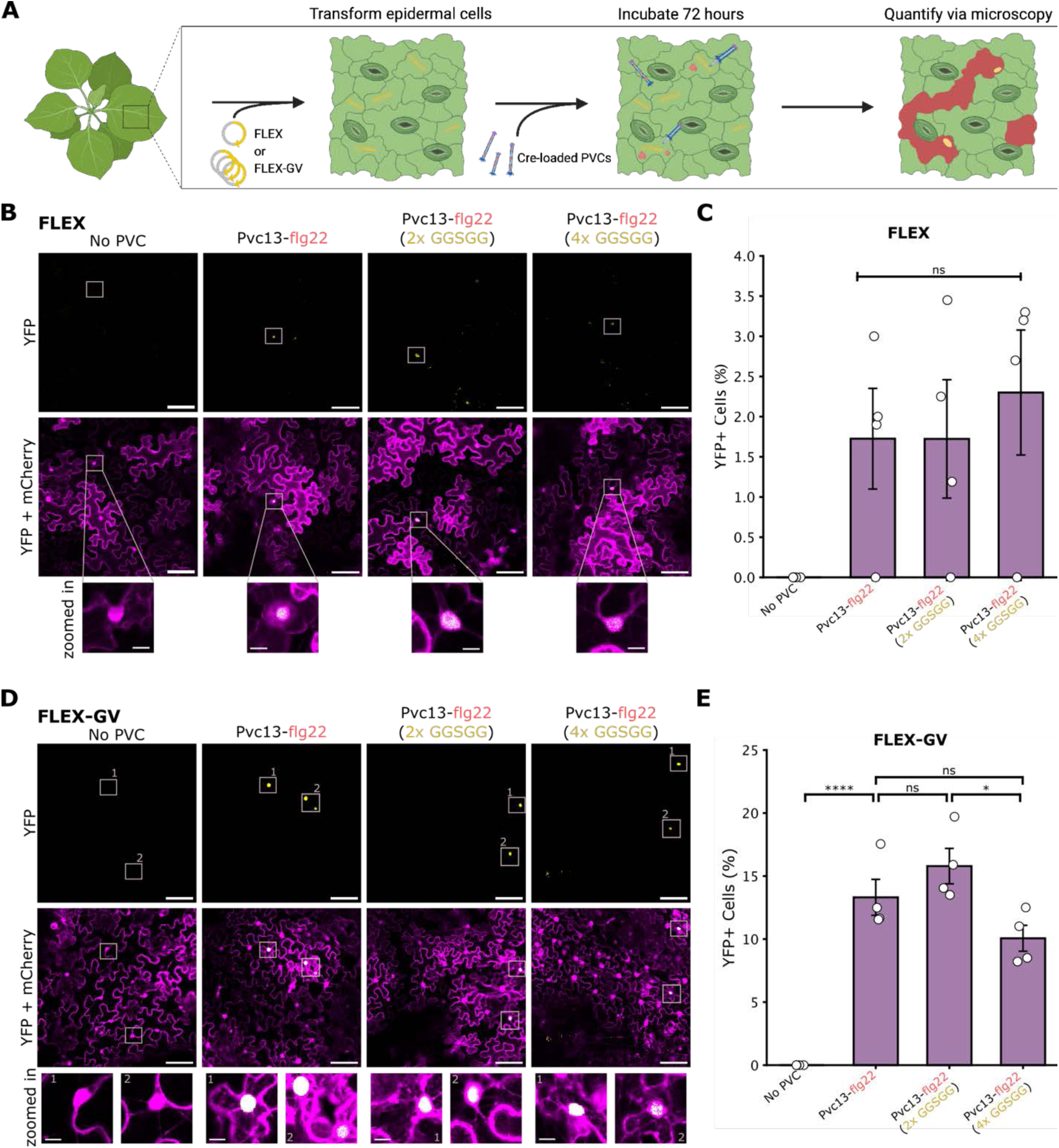
Reprogrammed PVCs deliver Cre recombinase to *N. benthamiana* leaves. **(A)** Schematic of the PVC delivery workflow in *N. benthamiana* leaves, where successful delivery events are registered as nuclear YFP. **(B)** Representative confocal fluorescence microscopy images of epidermal cells transformed with the FLEX reporter and treated with the Cre-loaded FLS2-targeting PVC library. Scale bar, 100 µm. Inset, 40 µm. **(C)** Quantification of the %YFP+ cells from confocal images in panel B using the FLEX reporter. **(D)** Representative confocal fluorescence microscopy images of epidermal cells transformed with the FLEX-GV reporter and treated with the Cre-loaded FLS2-targeting PVC library. Scale bar, 100 µm. Inset, 40 µm. **(E)** Quantification of %YFP+ cells from confocal images in panel D using the FLEX-GV reporter. Data are mean ± SEM with n = 4 biological replicates; one-way ANOVA with Tukey post hoc test. *p < 0.05, and ****p < 0.0001. ns, not significant.

We validated PVC-mediated Cre delivery and activity at the molecular level by sequencing FLEX-GV junctions following delivery of Pvc13-flg22 (2×GGSGG). Sequence alignments at the 5′ *lox* junction showed preferential retention of the *lox2272* site after inversion-excision, consistent with successful Cre-mediated recombination (**Fig. S10**). Inverted reads were detected in all Cre-treated samples, including those treated with Pvc13-flg22 nanoparticles, and were benchmarked against an *Agro*-delivered Cre construct control (**Fig. S10**). Together, these results demonstrate PVC-mediated delivery of functional Cre protein to intact, walled plant cells.

### PVC protein delivery is receptor-mediated and scales with receptor abundance

PVC binding and delivery depend on sufficient receptor density on the host cell membrane;^52^ therefore, we asked whether increasing FLS2 abundance would enhance delivery (**Fig. 5A**). First, we transiently overexpressed GFP-tagged FLS2 in *N. benthamiana* leaves via *Agro*-infiltration. The native *FLS2* promoter drove an approximately 8-fold increase in total FLS2 abundance relative to wild-type leaves (**Fig. S11A, B**), and most overexpressed FLS2-GFP colocalized with a fluorescent plasma-membrane marker (**Fig. 5B, Fig. S11C**).

**Figure 5.**
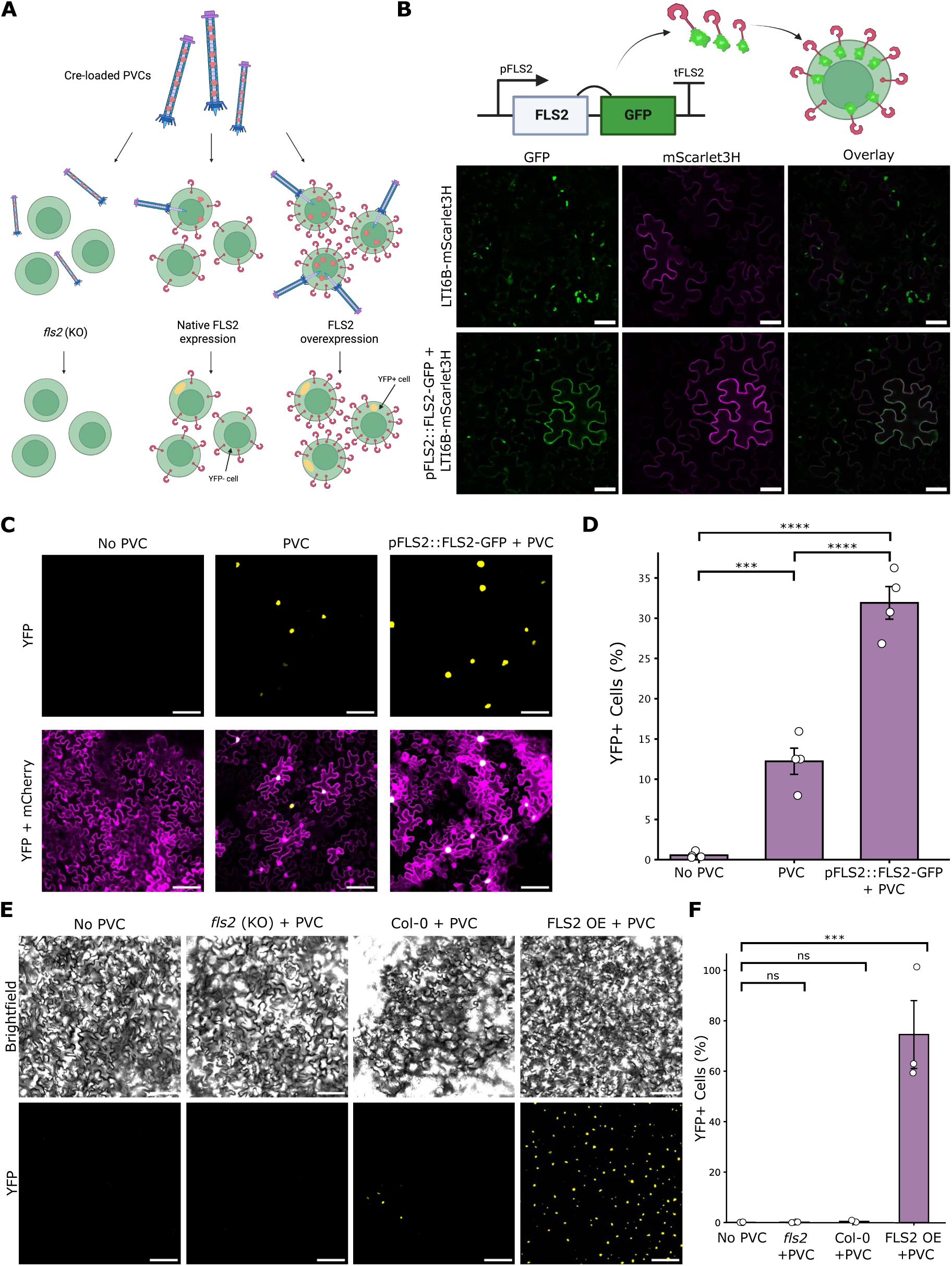
Reprogrammed PVC delivery is dependent on target receptor density. **(A)** Schematic of an activity assay for engineered PVCs based on receptor cell surface density. *fls2* knockout (KO) lines provide cells without the target receptor, and stable and transient overexpression (OE) of FLS2 provides cells with increased receptor surface densities. **(B)** FLS2 OE can be monitored by a GFP tag. Confocal microscopy of *N. benthamiana* leaves transiently co-expressing FLS2-GFP constructs and the plasma membrane marker LTI6B-mScarlet3H at 4 days post-infiltration (pFLS2::FLS2-GFP). Scale bar, 50 µm. **(C)** Representative confocal microscopy images of transient FLS2 OE in *N. benthamiana* leaves challenged with Cre-loaded Pvc13-flg22 (2xGGSGG). No PVC and no OE are included as controls. Scale bar, 100 µm. **(D)** Quantification of %YFP+ cells from confocal images in panel C. **(E)** Representative confocal microscopy images of *fls2* (KO) and FLS2 OE stable *A. thaliana* leaves challenged with Cre-loaded Pvc13-flg22 (2xGGSGG). No PVC and no OE are included as controls. Scale bar, 100 µm. **(F)** Quantification of %YFP+ cells from confocal images in panel E. Data in D and F are mean ± SEM with n = 3 biological replicates; one-way ANOVA with Tukey post hoc test. ***p < 0.001, and ****p < 0.0001. ns, not significant.

When Pvc13-flg22 (2×GGSGG) nanoparticles were applied to *N. benthamiana* leaves transiently overexpressing FLS2, Cre-driven YFP expression became more frequent and brighter (**Fig. 5C**). FLS2 overexpression increased delivery efficiency to an average of 32%, a more than 2.5-fold increase over delivery efficiency with the basal FLS2 expression (**Fig. 5D**). Molecular detection of FLEX inversion-excision events via NGS confirmed increased Cre activity after transient FLS2 overexpression (**Fig. S10C, F**).

To further test receptor dependence, we applied Cre-loaded Pvc13-flg22 (2×GGSGG) particles to stable *Arabidopsis thaliana fls2* knockout and FLS2-overexpression lines. Cre delivery was fully abolished in *fls2* knockout plants, confirming that FLS2 is required for PVC-mediated delivery (**Fig. 5E, F**). Conversely, stable FLS2-overexpression plants showed significantly increased Cre delivery, with efficiencies exceeding 75%. To note, delivery to wild-type *A. thaliana* leaves was lower than to wild-type *N. benthamiana* leaves.

We also used stable *Arabidopsis thaliana* pFRK1::mVenus reporter lines to assess activation of FLS2-dependent downstream signaling. Free flg22 induced strong mVenus expression, as expected, and insect-targeting PVCs produced detectable reporter activation, likely reflecting residual flagellin-derived activity in the bacterial preparation. However, FLS2-targeting PVC variants induced substantially higher mVenus expression than insect-targeting PVCs and were accompanied by altered expression of downstream FLS2-responsive genes *RBOHD* and *TIR1* (**Fig. S12**). These results support engagement of the FLS2 signaling pathway by FLS2-targeting PVCs.

### PVCs enable targeted chromosomal insertion in plants via R2 editor delivery

We next tested whether PVCs could deliver additional genome-engineering proteins using the R2 retrotransposon-derived genome editor, which enables site-specific DNA insertion into 25S ribosomal DNA (rDNA) safe-harbor loci in plants^53^. We engineered the R2 protein for PVC loading using the cargo-tagging strategy described above. After delivery into the cytosol, R2 forms a ribonucleoprotein complex with payload mRNA transcribed from a co-delivered payload plasmid, catalyzing reverse transcription and chromosomal integration of the payload (**Fig. 6A**). Following *Agro*-transformation of *N. benthamiana* leaves with an mCherry payload plasmid, we challenged them with R2-loaded Pvc13-flg22 (2×GGSGG) nanoparticles and quantified integration efficiency by droplet digital PCR (ddPCR) targeting the expected 3′ junction at the 25S rDNA locus, supplemented by sequence-level validation by deep NGS (**Fig. 6B**).

**Figure 6.**
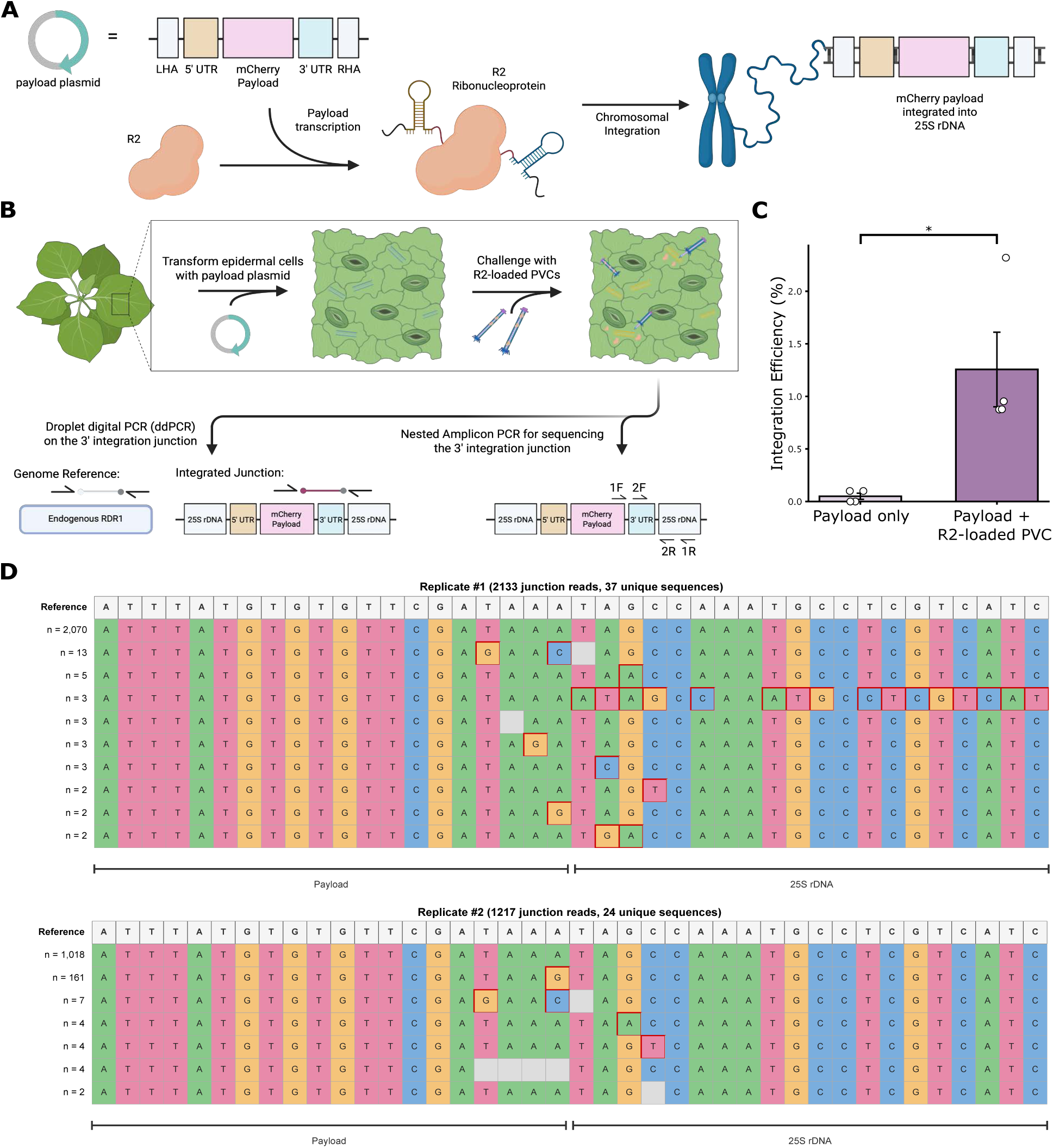
PVCs can deliver the R2 retroelement for targeted genome insertion in plants. **(A)** R2 ribonucleoprotein formation between a transcribed payload and an exogenously delivered R2 retroelement enables insertion of the payload into 25S rDNA safe harbor sites. **(B)** Schematic demonstrating the use of a payload plasmid as a reporter for PVC-mediated R2 delivery, in which R2 ribonucleoprotein activity is detected via ddPCR for quantification and NGS to confirm insertion events. **(C)** Integration efficiency of the R2 payload into *N. benthamiana* leaf cells, as detected by ddPCR, calculated as the percentage of payload integrations detected relative to the single-copy endogenous marker gene *NbRDR1*. PVC application was 1 mg mL^-1^ Pvc13-flg22 (2xGGSGG) loaded with R2. **(D)** Multiple sequence alignments of two biological replicates from panel C, following nested PCR to detect the 3’ integration junction. Data in C are mean ± SEM with n = 4 biological replicates; one-way ANOVA with Tukey post hoc test. *p < 0.05.

R2-loaded PVCs achieved payload integration efficiencies of approximately 1.3%, defined as the percentage of detected payload integrations relative to the single-copy endogenous marker gene *NbRDR1*. In contrast, payload-only controls produced negligible insertion (**Fig. 6C**). Sequencing of the 3′ integration junction confirmed mCherry payload insertion at the expected rDNA safe-harbor locus, with canonical junction sequences detected in the majority of reads across two sequenced biological replicates (**Fig. 6D**). These results demonstrate PVC-mediated delivery of a genome-engineering protein for targeted chromosomal insertion in plants.

Finally, we evaluated PVC cargo-size tolerance using a cargo library spanning the molecular weights of 79 kDa to 333 kDa, including multi-protein fusions of mScarlet, Cre, Cas9, and R2. All cargoes were loaded at comparable efficiency, as shown by HiBiT quantification, with no apparent size-dependent loading bias (**Fig. S13**), indicating that PVCs can accommodate large effector proteins relevant to plant genome engineering.

## Discussion

Here, we engineered PVC eCIS to target a natural plant cell-surface receptor and deliver proteins into plant cells. We targeted FLS2 as an initial proof-of-concept receptor and showed that reprogrammed PVCs deliver protein cargoes to both protoplasts and walled leaf cells, with efficiencies exceeding 75% in FLS2-overexpressing lines. To tune epitope accessibility, we designed PVC variants with (GGSGG)_n=0,2,4_ linkers flanking the tail-fiber-displayed flg22 epitope. Delivery efficiencies were similar across linker lengths, suggesting that epitope accessibility may not be a dominant limitation under the tested conditions. Engineered PVCs required different application conditions in protoplasts and leaf tissue. In protoplasts, Cre reporter activity peaked within 24 h, whereas in leaves it plateaued after 3 days. This slower delivery in leaf tissue could reflect reduced PVC diffusion and receptor access across the cell wall, as well as differences in reporter delivery, accumulation, and expression between the two systems. Future work should study how PVCs traverse the cell wall across different plant species, tissues, developmental stages, and receptor targets, as wall architecture and receptor accessibility may become limiting factors.

More broadly, this work introduces receptor-mediated delivery as a design principle for plant protein delivery. Receptor targeting is foundational in mammalian delivery, where ligand-receptor interactions are routinely exploited to improve cellular uptake, tissue tropism, and delivery specificity^54–57^. In plants, by contrast, delivery technologies have largely relied on physical disruption, passive uptake, nonspecific nanoparticle interactions, or bulk tissue transformation, lacking the ability to program entry via defined cell-surface receptors. Retargeted PVCs address this gap by coupling engineered receptor recognition to a contractile injection mechanism. This separates the delivery problem into modular and engineerable components: receptor choice can define tropism, tail-fiber design can tune binding specificity and signaling consequences, and cargo engineering can determine biological function. With further development, this approach could shift plant delivery from largely tissue-level or passive methods toward targeted, cell-specific delivery strategies analogous to those that have transformed delivery in mammalian systems.

The potential for cell-type-specific PVC delivery in plants remains unexplored. In mammalian systems, receptor-targeted PVCs can recognize cognate receptors with minimal off-target^44^. Our results establish FLS2 targeting as a proof of concept in *Arabidopsis thaliana* and *Nicotiana benthamiana*, but broader generality will require retargeting PVCs to additional receptors, tissues, developmental stages, and plant species. Targeting of cell-type-enriched receptors could reduce unintended delivery and provide a route toward spatially restricted cargo activity. Future development of PVCs targeting multiple plant receptors could also enable parallel delivery of distinct cargoes to different cell types or tissues, thereby expanding programmability. Alternatively, heterologous receptors orthogonal to plant receptors could be introduced as defined PVC targets, allowing delivery efficiency and specificity to be tuned for particular applications.

Microbe-associated molecular pattern receptors are central regulators of plant immune responses to pathogens^58^, and FLS2 is among the best-characterized examples. Its high surface abundance and well-characterized peptide binder, flg22, make FLS2 a practical first target for PVC retargeting in plants, but PVC-flg22 binding can activate immune signaling and receptor recycling^49^. Although we have not detected any detrimental effect on plant health in our study, the effects of this activation and repeated or long-term PVC delivery will require further study. Future strategies could mitigate this limitation by using flg22 variants that retain FLS2 binding but reduce or antagonize immune activation^59^, or by identifying allosteric or *de novo* binders to FLS2 or other plant receptors that support PVC attachment without perturbing native receptor signaling.

Finally, we demonstrated the utility and modularity of PVCs by delivering Cre recombinase (∼52 kDa) and by extending the platform to the larger R2 retroelement genome editor (∼147 kDa), thereby enabling targeted gene insertion. For genome-engineering applications, an important next step will be testing PVC delivery in regenerable plant materials, including callus, meristems, reproductive tissues, and crop transformation pipelines. Because receptor abundance and accessibility likely vary across tissues and developmental stages, future PVC designs should align receptor targets with the biological context necessary for the stable recovery of edited plants.

Although our R2 experiments demonstrated PVC-mediated delivery of a genome-engineering protein and targeted chromosomal insertion, the current assay still uses a plasmid-encoded DNA payload. Fully DNA-free genome engineering will require delivery of all editing components as proteins, RNAs, or ribonucleoprotein complexes. This could be achieved by engineering PVCs to co-deliver gene-editing proteins with guide or template RNAs^45^, or by using genome editors that do not require nucleic acids for editing, such as zinc-finger nucleases, TALENs, or guide-independent genome editors. Together, our results establish PVC eCIS as receptor-mediated protein-delivery vehicles for plant cells for the first time and support their further development for DNA-free genome-engineering applications in plant biotechnology and agricultural engineering.

## Methods

### Plant growth conditions

*Nicotiana benthamiana* and *Arabidopsis thaliana* (Col-0) plants were grown in a controlled-environment growth chamber (Conviron) set to 22°C during the day and 20°C during the night (16 h:8 h light/dark photoperiod) at 55% relative humidity.

### Plasmid construction

The PVCpnf gene locus had previously been synthesized and domesticated to produce the pStructural (Addgene pAWP78-PVCpnf1-16) and pCargo (Addgene pBR322-PVCpnf17-22). To generate a version of pCargo compatible with Golden Gate Assembly with BsaI, a single BsaI site was removed from the original pCargo construct using the Q5 Site-Directed Mutagenesis Kit (NEB E0554S). All further manipulations of either construct involved standard PCR amplification with Phusion High-Fidelity PCR Master Mix with HF Buffer (Thermo Fisher F531) followed by Golden Gate Assembly with BsaI HFv2 (NEB R3733) and T4 DNA Ligase (Thermo Fisher EL0014). Assembled constructs were transformed into chemically competent NEB Turbo cells (NEB C2984). For PVC production, final variants of pStructural and pCargo plasmids were co-transformed into chemically competent EPI300 cells (Fisher Scientific NC1583291).

### PVC purification

To generate a given PVC condition, one structural variant and one cargo variant were co-transformed into chemically competent EPI300 cells. The resulting transformants were then cultured, and PVC particles were harvested using a modified version of a previously developed method^48^. In brief, colonies were grown overnight in 2xYT (Thermo Fisher 22712020) medium and inoculated (at 1:1,000) into 1 L of Terrific Broth (Fisher Scientific BP246850), then shaken at 24°C for 48 h. Cultures were centrifuged at 4,000*g* for 20 min at 4°C and the resulting pellet was gently resuspended in 60 mL Buffer P (25 mM Tris-HCl pH 7.5 (Thermo Fisher 15567027), 140 mM NaCl (Sigma-Aldrich S5886), 3 mM KCl (Fisher Scientific P217), 5 mM MgCl_2_ (Sigma-Aldrich M2393), 200 μg mL^-1^ lysozyme (Thermo Fisher 89833), 50 μg mL^-1^ DNase I (Sigma-Aldrich DN25), 0.5% Triton X-100 (Sigma-Aldrich X100), and 1 Protease Inhibitor Cocktail (Sigma-Aldrich 11836153001) using a serological pipette and subsequently shaken at 250 rpm for 30 min at 37°C. Lysates were pelleted at 4,000*g* for 30 min at 4°C to remove cell lysate and the supernatant extracted and ultracentrifuged at 120,000*g* for 1 h at 4°C to pellet PVCs. The supernatant was discarded, and the ultracentrifuge tube was swabbed using a Kimwipe. PVC pellets were then washed with 1x PBS (Thermo Fisher 70011044), and the tubes were swabbed once more. Pellets were allowed to rehydrate overnight at 4°C in 2 mL of PBS before being resuspended by pipetting. Suspensions were agitated for 30 minutes to ensure complete rehydration, then centrifuged at 16,000*g* for 20 min at 4°C to clarify the solution. Supernatants were then diluted in 60 mL of PBS and ultracentrifuged at 120,000*g* for 1 h at 4°C once more. Tubes were swabbed after decanting the supernatant, and PVC pellets were resuspended by pipetting into 50 μL of working solution following a 4 h incubation period. Once again, suspensions were agitated for 30 min and then centrifuged at 16,000*g* for 20 min at 4°C to clarify the solution. The supernatant was collected as the final PVC product, and protein concentration was measured using a Qubit instrument (Thermo Fisher Q33211).

### In silico protein structure prediction

All protein structures and self-assemblies were predicted using ColabFold (v1.5.5), an AlphaFold2 implementation based on Google Colab that generates sequence alignments using MMseqs2. For general protein structure prediction, sequences were queried with the default model (AlphaFold2-ptm) and MSA settings. For Pvc13 tail fiber complex predictions, sequences were queried as homotrimers with the same default model (AlphaFold2-multimer-v3) and MSA settings. The resulting structures were rendered and customized with PyMOL Molecular Graphics System (v3.1.3).

### SDS-PAGE and denatured Western blotting

20 μg of purified PVCs was prepared for SDS-PAGE by combining with NuPAGE LDS Sample Buffer (ThermoFisher NP0007) and NuPAGE Sample Reducing Reagent (ThermoFisher NP0009) at appropriate dilutions. The mixture was then incubated for 10 min at 70°C in a thermocycler. The denatured PVC preparations were then loaded into NuPAGE Bis-Tris Mini Protein Gels, 4-12% (ThermoFisher NP0321) and run for 50 min at 200 V in 1x NuPAGE MOPS SDS Running Buffer (ThermoFisher NP0001) supplemented with NuPAGE Antioxidant (ThermoFisher NP0005) (at 1:400). For Coomassie staining, gels were rinsed of Running Buffer using Milli-Q ultrapure water prior to incubation in SimplyBlue SafeStain (ThermoFisher LC6065) for 90 min under gentle agitation. Gels were subsequently destained overnight in Milli-Q water and imaged in a Gel Imager (Azure Biosystems).

For western blot analysis of loaded PVC payloads, identical sample preparation and electrophoresis conditions were applied. In this case, PVCs were generated with loaded cargoes fused to a HiBiT peptide tag at the C-terminus, and unloaded cargoes were removed from the sample by standard ultracentrifugal separation. Variable quantities of PVC samples were applied to the above SDS-PAGE protocol, depending on the cargo loading capacity. Following electrophoresis, gels were blotted onto PVDF membranes (BIO-RAD 1704156) using a Trans-Blot Turbo Transfer System (BIO-RAD) set to default settings for turbo transfer of a mini gel (7-minute protocol). Finally, HiBiT-tagged cargo proteins were visualized using the Nano-Glo HiBiT Blotting System (Promega N2410), consisting of a 4 h TBST incubation and a 2 h LgBiT incubation, both at room temperature. Chemiluminescent images were captured with an Azure 200 Gel Imager. Band intensity analysis was performed using Fiji, a distribution of ImageJ.

For *in vitro* HiBiT detection of loaded proteins, the Nano-Glo HiBiT Lytic Detection System (Promega, catalog no. N3030) was used to denature PVC formulations and quantify HiBiT concentrations. In brief, 10 µg of a given PVC formulation was diluted to a final volume of 25 <u>µ</u>L in water and dispensed into an opaque, black 96-well plate (Costar). An equal volume of freshly prepared HiBiT Lytic reagent was added to the loaded wells, and the plate was immediately processed on a Tecan SPARK plate reader. The plate reader program allowed for 10 minutes of orbital shaking (magnitude = 2 mm) followed immediately by chemiluminescent detection (attenuation = None, integration time = 100 ms, settle time = 0 ms). All HiBiT detection assays were supplemented with a fresh standard curve using a HiBiT Control protein (Promega) within its linear range (10^-18^-10^-12^ moles). Samples were run as only a single technical replicate to avoid variation in chemiluminescent signal caused by small differences in incubation times.

### Electron microscopy

#### PVC formulations

300-mesh copper grids with Formvar/carbon support (PELCO®, Ted Pella, Inc., 01753-F) were glow-discharged in air for 1 min at 15 mA in negative-polarity mode. Purified PVC product suspended in PBS was diluted to 200 ng μL^-1^ in Milli-Q ultrapure water, and 5 µL was applied to the glow-discharged carbon film side of each grid for 1 min before side-blotting with filter paper. Grids were then washed twice by depositing 5 µL of Milli-Q water, followed by immediate blotting. Grids were stained with 5 µL of 1% (w/v) uranyl acetate in Milli-Q water for 30 s and blotted to remove excess stain. Prepared grids were examined on an FEI Tecnai T12 transmission electron microscope (120 kV, LaB₆ filament) equipped with a Gatan Ultrascan 2k × 2k CCD camera (Caltech Biological and Cryo-EM Facility). Images were acquired at nominal magnifications of 6,500×, 15,000×, and 30,000×.

#### Leaf tissue

Fully developed *Nicotiana benthamiana* leaves (the 3rd of 4th true leaf) from plants 3-4 weeks old were infiltrated with a high concentration (5 mg mL^-1^) of Pvc13-flg22 (2x GGSGG) nanoparticles. Leaf sections adjacent to the infiltration site were excised 24 hours post-infiltration and prefixed with 3% glutaraldehyde, 1% paraformaldehyde, 0.5% sucrose in PBS. Samples were further dissected with a scalpel under a dissecting microscope and transferred to brass planchettes (Ted Pella, Inc.) prefilled with an extracellular cryoprotectant buffer consisting of 10% Ficoll and 5% sucrose in 0.1 M sodium cacodylate trihydrate. Samples were rapidly frozen with a Wohlwend Compact-03 high-pressure freezing machine (Technotrade International, Inc.). The frozen samples were transferred under liquid nitrogen to cryotubes (Nunc) containing a frozen solution of 2.5% osmium tetroxide, 0.05% uranyl acetate in acetone. Tubes were loaded into an AFS-2 freeze-substitution machine (Leica Microsystems, Vienna) and processed at −90°C for 72 h, warmed over 12 h to −20°C, held at that temperature for 6 h, then warmed to 4°C for 2 h. The fixative was removed, and the samples were rinsed 3 times with cold acetone, after which they were infiltrated with Epon-Araldite resin (Electron Microscopy Sciences, Port Washington, PA) for 48 h. The samples were then flat-embedded between Teflon-coated glass microscope slides, and the resin was polymerized at 60°C for 48 h.

Embedded leaf samples were sectioned (170 nm) using a diamond knife (Diatome Ltd, Switzerland). Sections were placed on Formvar-coated, copper-rhodium slot grids (Electron Microscopy Sciences) and stained with 3% uranyl acetate and lead citrate. Grids were placed in a dual-axis tomography holder (Model 2040, E.A. Fischione Instruments, Export, PA) and imaged with a Tecnai T12-G2 transmission electron microscope operating at 120 KeV (Thermo Fisher Scientific) equipped with a 2k x 2k CCD camera (XP1000; Gatan, Inc.). Tomographic tilt series and large-area montaged overviews were acquired automatically using the SerialEM software package^60^. For tomography, samples were tilted ±62° and images collected at 1° intervals. The grid was then rotated 90°, and a similar series was taken about the orthogonal axis. Tomographic data were calculated, analyzed, and modeled using the IMOD software package^61–63^ on Mac Studio M1 and M3 computers.

#### Circular Dichroism

Spectral analysis was performed on PVC preparations within 1 day of nanoparticle isolation. CD spectra were acquired using an AVIV Circular Dichroism Spectrophotometer (Model 410). Far-UV spectra were recorded for 0.1 mg mL^-1^ PVCs suspended in pure Milli-Q water using a 1-mm path-length cuvette. Wavelength spectra were recorded at a constant temperature of 25 °C, and thermal melts were measured at 220 nm. Near-UV spectra were recorded for 1 mg mL^-1^ PVCs suspended in pure Milli-Q water using a 10 mm path-length cuvette. Wavelength spectra were recorded at a constant temperature of 25 °C, and thermal melts were recorded at 292 nm. Melting temperatures were extracted from thermal melts by fitting the data to a sigmoidal function and extracting the inflection point from the resulting curve.

#### Protoplast isolation

Protoplasts were isolated from 4-week-old *Arabidopsis thaliana* leaves as described previously^64^, with some modifications. In brief, 7-10 fully developed leaves were gently compressed between Time tape (adhered to the upper epidermis) and 3 M Magic tape (adhered to the lower epidermis). The lower epidermal layer of the leaves was removed and discarded, thus exposing the leaf mesophyll. The exposed mesophyll was then placed in contact with a cell wall-degrading enzyme solution (20 mM MES pH 5.7, 0.4 M mannitol, 20 mM KCl, 1.5% w/v cellulase R10 Yakult, and 0.4% w/v macerozyme R10 Yakult) in the dark and incubated for 3 h. Protoplasts released into the enzyme solution following incubation were diluted in ice-cold W5 solution (2 mM MES pH 5.7, 154 mM NaCl, 125 mM CaCl_2_, and 5 mM KCl) and gently pelleted at 100*g* for 5 min with minimal ramp rates. Pellets were suspended in W5 solution and run on a 21% w/v water-based sucrose cushion at 90*g* for 10 min. The suspended layer of intact protoplasts was once again diluted in W5 solution and gently pelleted at 100*g* for 5 min with minimal ramp rates. The final pellet was resuspended to 5×10^5^ cells mL^-1^ in working solution.

#### Plasmid transfection for protoplast assays

For assays requiring transfection of a FLEX reporter plasmid and/or a Cre expression plasmid, DNA was transfected into protoplasts using a PEG-mediated transfection procedure. Briefly, 100 μL of protoplasts at 5×10^5^ cells mL^-1^ in an MMG working solution (4 mM MES pH 5.7, 0.4 M mannitol, and 15 mM MgCl_2_) were gently mixed with 40 μg total plasmid DNA (20 μg:20 μg co-transfections) and 100 μL PEG transfection solution (40% w/v PEG4000, 0.2 M mannitol, 0.1 M CaCl_2_) in a round-bottom tube. After 15 min of incubation, the protoplasts were washed with 1 mL W5 solution and pelleted at 200*g* for 2 min. This wash step was repeated three times, and the final protoplast pellet was resuspended in WI solution (4 mM MES pH 5.7, 0.5 M mannitol, and 20 mM KCl) at a density of 10^5^ cells mL^-1^.

#### PVC delivery to protoplasts

All protoplast delivery experiments were performed using fresh *Arabidopsis thaliana* mesophyll protoplasts isolated using the aforementioned procedure. Protoplasts in a working solution of WI media were seeded into clear-bottom 96-well plates (Fisher Scientific 07-000-167) and allowed to settle for 30 minutes prior to manipulation. For all experiments, 2×10^4^ cells were seeded into a given well, and excess WI solution was aspirated to a final volume of 50 μL. PVCs were then added to a final concentration of 500 ng μL^-1^, unless otherwise specified.

For those assays involving native toxin delivery, cells were incubated for 24 h following PVC injection before confocal analysis. For those assays involving fluorescent protein delivery, cells were incubated for 18 h (unless otherwise specified for dose-response analysis) following PVC injection of 1 mg mL^-1^ before confocal analysis. For those assays involving FLEX reporters and Cre protein delivery, cells were transfected with the appropriate FLEX reporter and seeded at 2×10^4^cells per well in WI working solution. Following PVC injection, cells were incubated for an additional 48 h (unless otherwise specified) before confocal analysis.

#### Quantification of PVC activity in protoplasts

For each PVC condition, 3 biological replicates (3 separate wells individually challenged with PVCs) were performed; for each biological replicate, 10 technical replicates (10 non-overlapping confocal fields of view) were collected. Images were taken around the perimeter of the wells due to the tendency of protoplasts to cluster along the well edges. Each field of view was analyzed with Fiji, a distribution of ImageJ, for quantification.

For those assays involving toxin delivery, cytotoxicity was measured using PI/FDA staining. Briefly, following a 24 h incubation with PVCs, cells were simultaneously stained with 20 ng mL^-1^ fluorescein diacetate (Invitrogen, F1303) and 400 ng mL^-1^ propidium iodide (Sigma-Aldrich, P4170). Cells were incubated for 10 min and then imaged under a Leica STELLARIS 8 FALCON confocal microscope. PI fluorescence was captured using a 535 nm laser excitation, and FITC fluorescence was captured using a 495 nm laser excitation from a white light laser. Images were rendered and analyzed using Fiji software. Cytotoxicity for a biological replicate was calculated by combining PI and FITC counts for all technical replicates and calculating Cytotoxicity = (# PI+ cells)/(# PI+ cells + # FITC+ cells).

For those assays involving fluorescent protein delivery, cells were imaged under a Leica STELLARIS 8 FALCON confocal microscope. mScarlet fluorescence was captured using a 569 nm laser excitation from a white light laser. Images were rendered using FIJI and analyzed using a custom Python script. In each field of view, fluorescent nuclei were counted manually, and total cell counts were determined via Cellpose segmentation. mScarlet delivery for a biological replicate was calculated by combining mScarlet and total cell counts across all technical replicates and setting mScarlet delivery = (# mScarlet+ cells)/(# total cells).

For those assays involving FLEX reporters and Cre protein delivery, cells were imaged using a Leica STELLARIS 8 FALCON confocal microscope with 434 nm, 517 nm, and 587 nm laser excitations from a white-light laser to capture mTurquoise2, YPET, and mCherry expression, respectively. Images were rendered using FIJI and analyzed using a custom Python script. In each field of view, YFP+ nuclei were counted manually, and mCherry+ cells were determined via Cellpose segmentation. The PVC activity for a biological replicate was then calculated by combining YFP and mCherry counts across all technical replicates and defining PVC activity as (# YFP+ cells)/(# mCherry+ cells).

#### PVC delivery to leaf cells

For studies in *Nicotiana benthamiana*, fully developed leaves (the 3rd of 4th true leaf) from plants 3-4 weeks old were selected for all experiments. For studies in *Arabidopsis thaliana,* fully developed leaves (the 3^rd^ rosette) from plants 3-4 weeks old were selected for all experiments. Biological replicates were sampled from leaves on separate plants generated from the same seed batch. *Agrobacterium tumefaciens* strain GV3101 carrying the desired reporter plasmid, the p19 silencing-suppressor plasmid, and the pSOUP helper plasmid was cultured in 2xYT media (ThermoFisher22712020) supplemented with 10 μg mL^-1^ rifampicin (Sigma-Aldrich R3501), 20 μg mL^-1^ gentamycin (Sigma-Aldrich G1264), 50 μg mL-1 tetracycline (Sigma-Aldrich T7660), and 50 μg mL^-1^ kanamycin (Sigma-Aldrich K1637) at 30°C and 200 rpm for 24 h. Overnight cultures were then centrifuged at 4,000g for 20 min, and the pellets were resuspended to an OD600 of 0.5 in infiltration buffer (10 mM MES, pH 5.7 (Sigma-Aldrich M2933), 10 mM MgCl2 (Sigma-Aldrich M2393), 200 μM acetosyringone (PlantMedia 40100297). The resuspended cultures were incubated at room temperature and shaken at 120 rpm for 4 h in the dark. Following incubation, 100-200 μL of the *Agrobacterium* mixture was infiltrated against the abaxial surface of a leaf with a 1 ml needleless syringe by applying gentle pressure.

For assays involving FLEX reporters and Cre protein delivery, plants were incubated at normal growth conditions for 24 hours to allow plasmid transformation. Purified Cre-loaded PVCs resuspended in 10 mM HEPES Buffer, pH 7.2 (Sigma-Aldrich H3375) at a concentration of 1 mg mL^-1^ were infiltrated into the same leaf area through the abaxial surface using a 1 mL needleless syringe. Plants were incubated at normal growth conditions for 3 days (unless otherwise specified for time-course studies) before confocal analysis and/or tissue collection.

For assays involving transient FLS2 overexpression, a pre-incubation step was included to introduce the FLS2 OE construct before *Agro*-transformation of the reporter constructs. FLS2 OE constructs were harbored within the same *Agrobacterium* strains as before. They were cultured, induced, and infiltrated into leaves in an identical procedure. Plants were then incubated under normal growth conditions for 3 days prior to *Agro*-infiltrating the appropriate reporter construct (in our case, a FLEX reporter) and proceeding with the above protocol as specified. For Cre-delivery studies, this would involve a 3-day pre-incubation following *Agro*-infiltration of the FLS OE cassette, a 1-day incubation following *Agro*-infiltration of the FLEX reporter, and a 3-day incubation following PVC challenge; the resulting experimental procedure thus comprises 3 separate leaf infiltrations.

For assays involving integration payloads and R2 protein delivery, plants were incubated at normal growth conditions for 24 hours to allow plasmid transformation. Purified R2-loaded PVCs resuspended in 10 mM HEPES Buffer, pH 7.2 (Sigma-Aldrich H3375) at a concentration of 1 mg mL^-1^ were infiltrated into the same leaf area through the abaxial surface using a 1 mL needleless syringe. Plants were incubated at normal growth conditions for 5 days before tissue collection.

#### Quantification of PVC activity in leaves

For assays involving FLEX reporters and Cre protein delivery, PVC-infiltrated leaves were prepared for confocal imaging 72 h post-infiltration. A leaf disk puncher (Fisher Scientific NC0769832) was used to extract a 0.25 in leaf section adjacent to the site of PVC infiltration. Leaf disks were then mounted between a glass slide and a coverslip of #1 thickness using water as the mounting medium. A Leica STELLARIS 8 FALCON confocal microscope was used to image the plant tissue with 434 nm, 517 nm, and 587 nm laser excitations from a white light laser to capture mTurquoise2, YPET, and mCherry expression, respectively. All images were obtained at 20x magnification with water as an immersion medium. For each PVC condition, 3-4 biological replicates (3-4 infiltrations into leaves of separate plants) were performed, and for each biological replicate, more than 8 technical replicates (8 non-overlapping confocal fields of view) were collected. Each field of view was rendered in FIJI and analyzed using a custom Python pipeline to quantify the total number of YFP-expressing nuclei and mCherry-expressing cells in that field of view. The PVC activity for a biological replicate was then calculated by combining YFP and mCherry counts across all technical replicates and defining PVC activity as (# YFP+ cells)/(# mCherry+ cells).

For assays involving integration payloads and R2 protein delivery, genomic DNA was extracted from PVC-infiltrated leaves 5 days post-infiltration. Samples were then processed for ddPCR and NGS analysis as described in those methods.

#### Nested PCR for amplicon sequencing

For detection of the FLEX 3’ inversion-excision junction, 100 ng of genomic DNA was amplified by nested PCR using Phusion High-Fidelity PCR Master Mix (New England Biolabs, catalog no. M0531) in 25 µl reactions. The resulting reaction solution containing an OUT product mixture consisting of both inverted and non-inverted amplicons was purified using the QIAquick PCR Purification Kit (Qiagen, catalog no. 28104). The DNA concentration of the purified reaction was measured on a NanoDrop One (Thermofisher), and 10 ng was carried into the nested round. The nested reaction was performed in two separate reactions using Phusion High-Fidelity PCR Master Mix, resulting in a non-inverted IN product of 928 bp and an inverted IN product of 579 bp. Both products from the nested round were resolved on a 1% agarose gel, and the expected 579 bp band from the inverted product was excised and purified with the QIAquick Gel Extraction Kit (Qiagen). The purified nested inverted amplicons from two biological replicates were sequenced by Plasmidsaurus amplicon sequencing.

For detection of the R2 3′ integration junction, genomic DNA was amplified by nested PCR using Phusion High-Fidelity PCR Master Mix (New England Biolabs, catalog no. M0531) in 30 µl reactions. The resulting OUT product was 425 bp. The completed reaction was diluted 50-fold in water, and 2 µl of the dilution was carried into the nested round. The nested reaction was performed using Phusion High-Fidelity PCR Master Mix and resulted in an IN product of 215 bp. Products from the nested round were resolved on a 1.5% agarose gel, and the expected 215 bp band was excised and purified with the QIAquick Gel Extraction Kit (Qiagen). Purified nested amplicons from two biological replicates were sequenced by Plasmidsaurus amplicon sequencing. Primer sequences used are listed in **Supplementary Table 3.**

#### Amplicon sequencing

PCR amplicons were submitted to Plasmidsaurus for Oxford Nanopore Technology (ONT) sequencing, yielding approximately 5,400-5,500 reads per sample in FASTQ format. Reads were filtered to retain only those containing specified flanking anchor sequences found in **Supplementary Table 3.** The intervening sequence between anchors was extracted and evaluated against the canonical reference sequence: for the R2 3’ integration junction (5’-TAAATAGC-3’) and for the lox2272-retained FLEX inversion-excision junction (5’-TATTTACAATTACAGCGGCCGCCCCGGAATAACTTCGTATAGGATACTTTATACGAAGTTATCCCGCCAAAAAAATGGCTGCAGCCAAGGGCGA-3’). For visualization, sequences were gap-aligned to the canonical reference sequence using difflib.SequenceMatcher. Aligned sequences were rendered as per-row character grids in matplotlib, with substituted positions outlined in red and deletion gaps represented as empty cells.

#### Membrane receptor immunoblotting

25 mg of *N. benthamiana* leaf tissue was subjected to two cycles of liquid nitrogen flash freezing and bead beating for 30 seconds using a Bead Mill 4 Homogenizer (Fisher Scientific 15-340-164). Homogenized samples were briefly centrifuged and mixed with 75 μL of 2x Laemmli SDS sample buffer (Serva 42526.01) supplemented with 4% (v/v) 2-mercaptoethanol (Gibco 21985-023). Samples were incubated for 10 min at 65°C, chilled on ice for 2 min, and clarified by centrifugation at 16,000 × g for 10 min at 4°C. Clarified supernatant was loaded at 10 μL per lane into a 4-20% Mini-PROTEAN TGX Precast Protein Gel (BIO-RAD 4561094). Following electrophoresis, proteins were blotted onto pre-assembled 0.2 μm nitrocellulose membrane transfer stacks (BIO-RAD 1704158) using a Trans-Blot Turbo system (BIO-RAD 1704150) set to the high-molecular-weight transfer protocol.

Following transfer, blots were rinsed with 1x TBS-T (20 mM Tris, 150 mM NaCl, pH 7.6) and blocked in either 5% non-fat milk powder in TBS-T for actin detection or 10% non-fat milk and 5% BSA in TBS-T for FLS2 detection. Membranes were incubated overnight at 4°C under gentle agitation with anti-β-actin antibody (1:1,000; Abcam ab197345) or anti-FLS2 antibody (1:1,000; PhytoAB PHY2628A) diluted in the respective blocking buffers. Membranes were then washed four times for 10 min each with 1x TBS-T and incubated for 1 h at room temperature with Goat Anti-Rabbit HRP-conjugated secondary antibody (1:10,000; BIO-RAD 1705046). Membranes were then washed four times for 10 min each with 1x TBS-T. The actin membrane was developed with Pierce ECL Western Blotting Substrate (Thermo Scientific 32106), and the FLS2 region of interest was developed with SuperSignal West Atto Ultimate Sensitivity Substrate (Thermo Scientific A38554). Band intensity analysis was performed using Fiji.

#### Confocal of membrane colocalization

Fully developed leaves from *Nicotiana benthamiana* plants were selected for all experiments. Two separate cultures of *Agrobacterium tumefaciens* (GV3101) harboring either the transient membrane marker LTI6B-mScarlet3H or a transient FLS2 expression cassette were grown in 2xYT media (ThermoFisher 22712020) supplemented with 10 μg mL^-1^ rifampicin (Sigma-Aldrich R3501), 20 μg mL^-1^ gentamycin (Sigma-Aldrich G1264), 50 μg mL^-1^ tetracycline (Sigma-Aldrich T7660), and 50 μg mL^-1^ kanamycin (Sigma-Aldrich K1637) at 30°C and 200 rpm for 24 h. Overnight cultures were then centrifuged at 4,000*g* for 20 min, and the pellets were separately resuspended to an OD600 of 0.5 in infiltration buffer (10 mM MES, pH 5.7 (Sigma-Aldrich M2933), 10 mM MgCl_2_ (Sigma-Aldrich M2393), 200 μM acetosyringone (PlantMedia 40100297). The resuspended cultures were incubated at room temperature and shaken at 120 rpm for 4 h in the dark. Following incubation, 100-200 μL of the *Agrobacterium* mixture was infiltrated against the abaxial surface of a leaf with a 1 ml needleless syringe by applying gentle pressure. For co-infiltration of LTI6B-mScarlet3H and the FLS2 expression cassette, the induced *Agro* cultures were combined at a 1:1 (v/v) ratio, resulting in a co-infiltration solution at OD 0.5. Leaves were incubated for 4 days prior to imaging on a Leica STELLARIS 8 FALCON confocal microscope. Leaf tissue was collected after this time point and flash-frozen in liquid nitrogen for downstream processing in membrane receptor studies.

#### pFRK1::mVenus reporter detection

*Arabidopsis thaliana* leaves of the reporter line were challenged with free flg22 peptide or engineered PVCs in HEPES buffer via syringe infiltration. Following 12 hours of incubation, a leaf disk puncher (Fisher Scientific NC0769832) was used to extract a 0.25 in leaf section adjacent to the site of PVC infiltration. Leaf disks were then mounted between a glass slide and a coverslip of #1 thickness using water as the mounting medium. A Leica STELLARIS 8 FALCON confocal microscope was used to image the plant tissue with a 515 nm laser excitation from a white light laser to capture mVenus expression. All images were obtained at 20x magnification with water as an immersion medium. For each PVC condition, 3-4 biological replicates (3-4 infiltrations into leaves of separate plants) were performed, and for each biological replicate, more than 8 technical replicates (8 non-overlapping confocal fields of view) were collected. Each field of view was rendered and analyzed in Fiji, a distribution of ImageJ, to quantify the raw intensity density of the mVenus signal. The mean raw integrated density (RID) for a biological replicate was then calculated by averaging the RID across all technical replicates.

#### Quantitative PCR

*Arabidopsis thaliana* leaves were challenged with PVCs in HEPES buffer via syringe infiltration. After 6 h of incubation, leaves were flash-frozen for downstream analysis. Total RNA was extracted from approximately 100 mg of frozen leaf tissue using the RNeasy Plant Mini Kit (Qiagen) with on-column DNase I digestion (RNase-Free DNase Set, Qiagen). In brief, tissue was homogenized under liquid nitrogen by bead beating (2 × 30 s at speed 5), and lysates were prepared in Buffer RLC supplemented with β-mercaptoethanol (10 µL per mL buffer). Clarified lysate was obtained by centrifugation through QIAshredder spin columns (16,000 × g, 2 min). After the addition of 0.5 volumes of ethanol, the lysate was applied to RNeasy Mini spin columns. On-column DNase I digestion was performed by applying 80 µL of DNase I incubation mixture (10 µL DNase I stock solution in 70 µL Buffer RDD per sample) directly to the spin column membrane for 15 min at room temperature. RNA was eluted in 30 µL of RNase-free water and quantified using the Qubit RNA BR Assay Kit (Thermofisher, Q10210) on a Qubit 4 Fluorometer (ThermoFisher Scientific).

First-strand cDNA was synthesized from 1 µg of total RNA in 20 µL reactions using the iScript cDNA Synthesis Kit (Bio-Rad). cDNA was diluted fivefold prior to use. RT-qPCR was performed on a CFX96 Touch System (Bio-Rad) using PowerUp SYBR Green Master Mix (ThermoFisher Scientific, A25742) in 10 µL reactions containing 12.5 ng cDNA, 1× master mix, and 750 nM of each primer pair (equimolar forward and reverse, premixed). Oligonucleotides were synthesized by Integrated DNA Technologies and are listed in Supplementary Table [X]. Thermocycler conditions were [50°C for 2 min; 95°C for 2 min; 40 cycles of 95°C for 15 s, 60°C for 60 s]. Relative gene expression was calculated using the 2^−ΔΔCt^ method and normalized to the reference gene *SAND*. All conditions were analyzed with 4 biological replicates and 4 technical replicates for each target gene. Oligonucleotides were synthesized by Integrated DNA Technologies and are listed in **Supplementary Table 5**.

#### Digital droplet PCR

Genomic DNA was extracted from 100 mg of *N. benthamiana* leaf tissue collected from the leaf infiltration site using the DNeasy Plant Mini Kit (Qiagen) according to the manufacturer’s protocol. Each 22 µl duplex ddPCR reaction contained 11 µl of ddPCR Supermix for Probes (no dUTP; Bio-Rad, catalog no. 1863024), 900 nM of each forward and reverse primer for the target and reference loci, 250 nM FAM-labeled, ZEN/Iowa Black FQ double-quenched probe targeting the integrated R2 3′ junction, 250 nM HEX-labeled, ZEN/Iowa Black FQ double-quenched probe targeting the NbRDR1 reference gene, and 50 ng of gDNA. Oligonucleotides were synthesized by Integrated DNA Technologies and are listed in Supplementary Table 4. Each reaction was loaded with 70 µl of droplet-generation oil (Bio-Rad, catalog no. 1863005) into a DG8 cartridge (Bio-Rad, catalog no. 1864007) and partitioned into droplets using a QX200 Droplet Generator. Forty microliters of the resulting emulsion were transferred to a 96-well plate, heat-sealed with pierceable foil, and thermally cycled per the manufacturer’s protocol with annealing/extension at 56°C. Droplets were read on a QX200 Droplet Reader and analyzed in QX Manager software (Bio-Rad). The average copy number of insertion per genome was calculated as the ratio of FAM-labeled R2 3′ junction copies to HEX-labeled NbRDR1 reference copies, assuming a single NbRDR1 copy per genome in *N. benthamiana*.

#### Statistics and reproducibility

Statistical analyses were performed using Prism (10.1.0). Quantitative data are presented as mean ± SEM with n = 3-4 biological replicates per condition. Figure legends provide further specification as necessary. Unless otherwise stated, biological replicates represent independent treatments in separate wells (*in vitro* assays) or on leaves from separate plants. All micrographs, gels, and blots are representative images from at least 2 independent repeated experiments. Statistical significance was computed using one-way or two-way ANOVA with Tukey post hoc tests (multiple comparisons correction), as indicated in figure legends. p < 0.05 was considered statistically significant.

## Data Availability Statement

All data are included either in the manuscript or in the supplementary files. All sequencing results are accessible through the National Center for Biotechnology Information Sequence Read Archive under BioProject <u>PRJNA1484212</u>^65^.

## Code Availability Statement

All code used in the study is available at https://github.com/DemirerLab. GraphPad Prism Version 10.3.1 and ImageJ were used to analyze all other data.

## Author Contributions

Conceptualization: MGL, GSD

Methodology: MGL, CAH, KTM, GSD

Data Acquisition: MGL, CAH, CC, KTM, YW, VP

Supervision: GSD

Writing: MGL, GSD Funding

Acquisition: GSD

## Acknowledgements

We thank the Nolan lab at Caltech for the pFRK1:mVenus Arabidopsis lines, and Niko Geldner for FLS2 transgenic Arabidopsis lines. Fluorescence imaging was performed at the Biological Imaging Facility, with support from the Caltech Beckman Institute and the Arnold and Mabel Beckman Foundation. Electron microscopy imaging was performed at the Caltech Biological and Cryo-EM Facility, with staff support for collecting leaf tissue tomograms. Schematics were partially created with BioRender.com. All Bio-Rad ddPCR equipment was provided by the CLARITY, Optogenetics, and Vector Engineering Research (CLOVER) Center at Caltech.

