## Supplementary Figures for "Extracellular contractile injection systems for receptor-mediated protein delivery into plant cells"

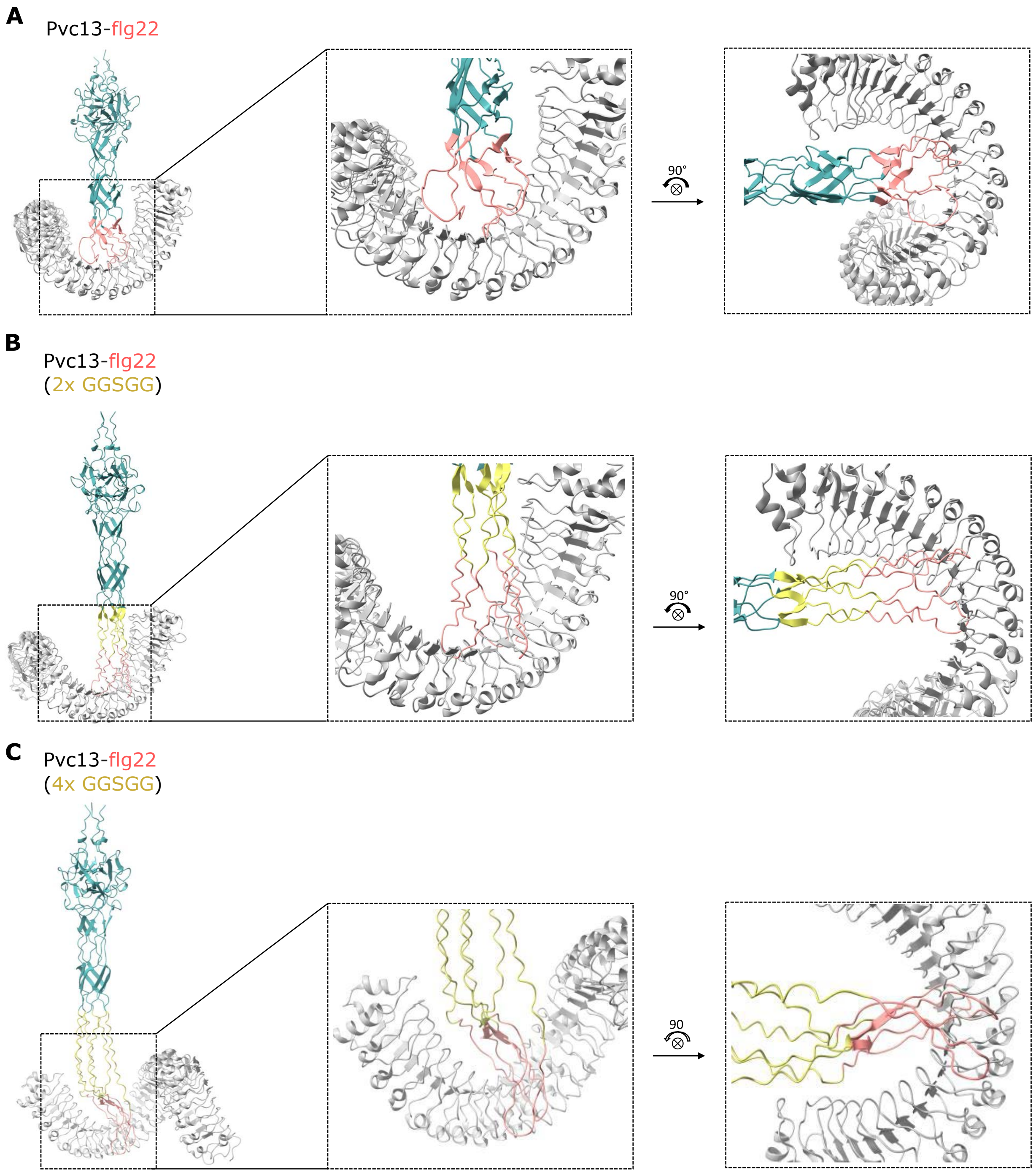


**Supplementary Figure 1. Pvc13-flg22 epitope alignments against the FLS2 receptor for all PVC variants. (A)** ChimeraX alignment of the flg22 epitope (salmon) from the Pvc13-flg22 PVC variant AlphaFold-predicted tail fiber body (teal) to the flg22 peptide in the FLS2–flg22 crystal structure (PDB: 4MN8, gray). **(B)** ChimeraX alignment of the flg22 epitope (salmon) from the Pvc13-flg22 (2xGGSGG) PVC variant AlphaFold-predicted structure (tail fiber body, teal; linker, yellow) to the flg22 peptide in the FLS2–flg22 crystal structure (gray). **(C)** ChimeraX alignment of the flg22 epitope (salmon) from the Pvc13-flg22 (4xGGSGG) PVC variant AlphaFold-predicted structure (tail fiber body, teal; linker, yellow) to the flg22 peptide in the FLS2–flg22 crystal structure (gray).

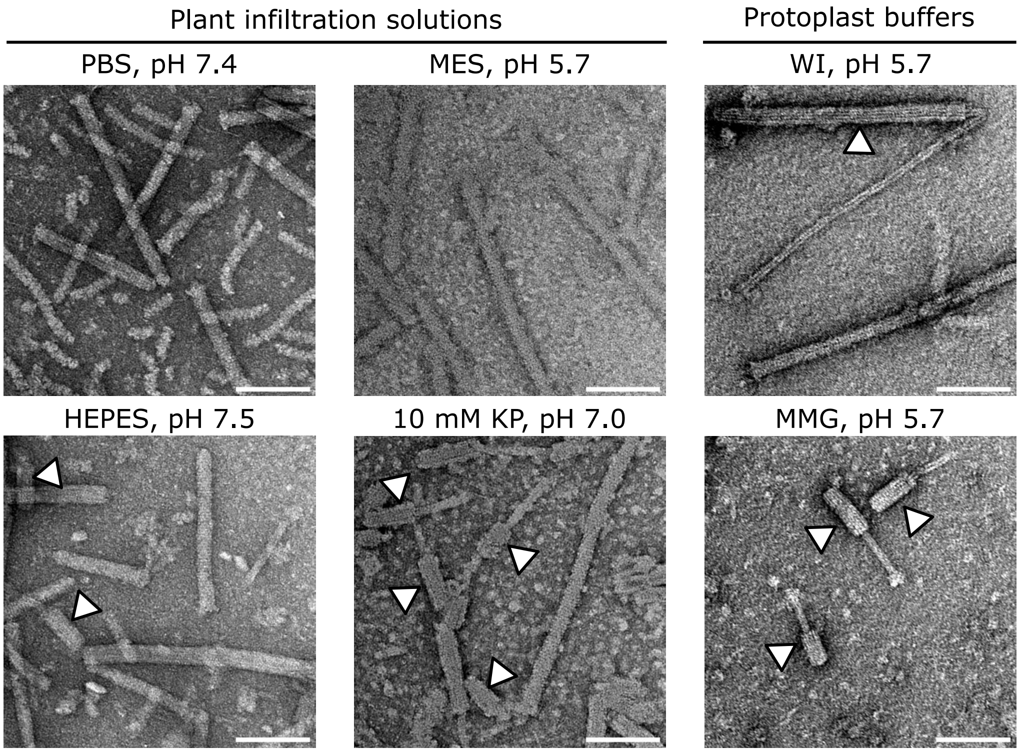


**Supplementary Figure 2. Characterizing PVC stability in different plant buffers.** Representative negative stain TEM images of Pvc13-flg22 nanoparticles prepared in various suspension buffers. White arrows indicate contracted PVC sheaths. Scale bars, 100 nm.

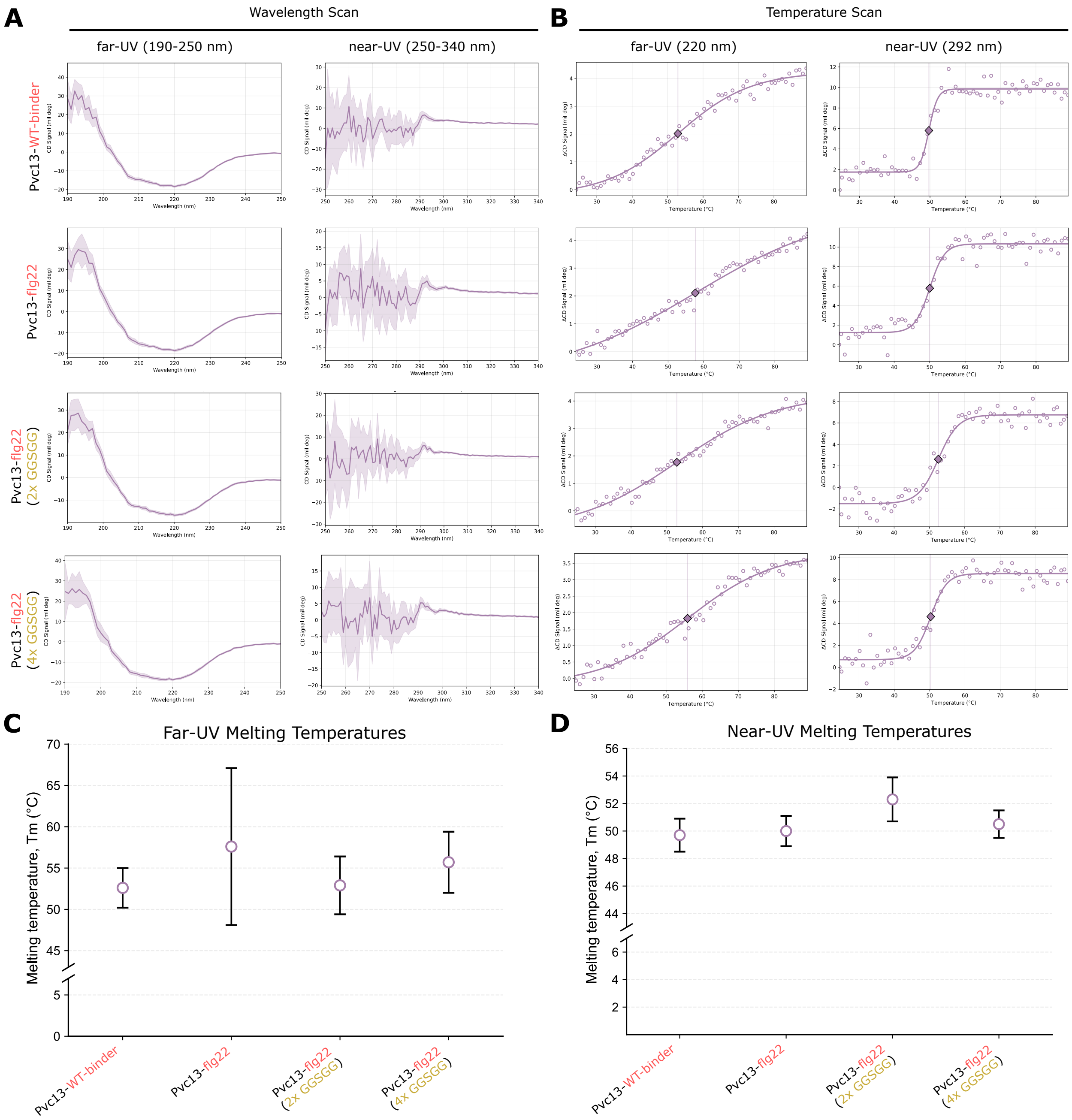


**Supplementary Figure 3. Tail fiber engineering does not alter the thermal stability of PVC complexes. (A)** Circular dichroism (CD) wavelength scans of all PVC variants in the far-UV (left) and near-UV (right) regions at 25 °C. **(B)** Circular dichroism melting curves for all PVC variants, measured at peak wavelengths in the far-UV (220 nm, left) and near-UV (292 nm, right) regions. Data are fitted to a sigmoidal function, and the inflection point of the sigmoidal fit is indicated by the dashed vertical line. **(C)** The far-UV melting temperatures of PVC variants, as collected from panel B, with errors propagated from the sigmoidal fit. **(D)** The near-UV melting temperatures of PVC variants, as collected from panel B, with errors propagated from the sigmoidal fit. Data represent a single biological replicate; error bars indicate the standard error of the fitted parameter. Differences in panels C and D are not statistically significant.

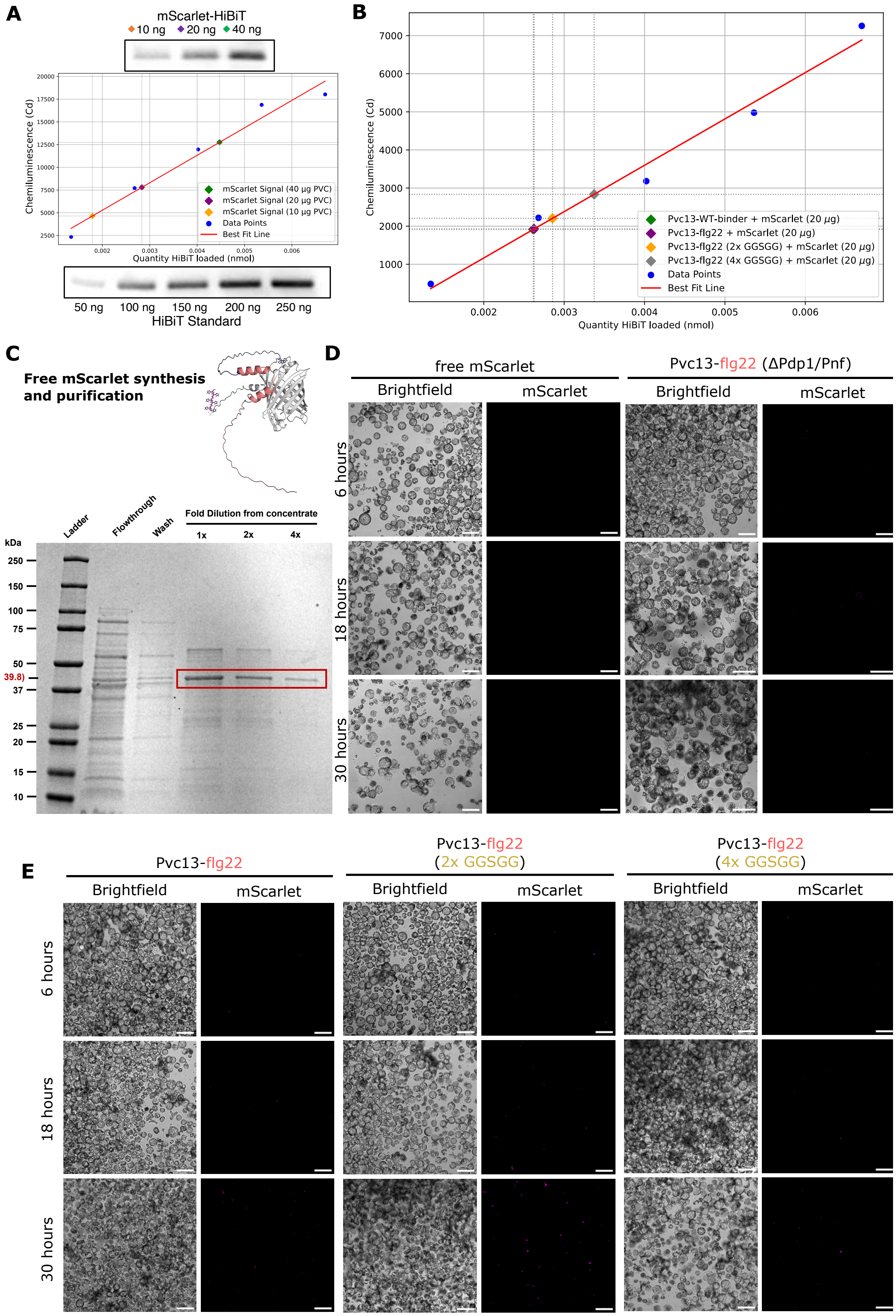
A
mScarlet-HiBiT
10 ng 20 ng 40 ng
Chemiluminescence (Cd)
Quantity HiBiT loaded (nmol)
mScarlet Signal (40 µg PVC)
mScarlet Signal (20 µg PVC)
mScarlet Signal (10 µg PVC)
Data Points
Best Fit Line
50 ng 100 ng 150 ng 200 ng 250 ng
HiBiT Standard
B
Chemiluminescence (Cd)
Quantity HiBiT loaded (nmol)
Pvc13-WT-binder + mScarlet (20 µg)
Pvc13-flg22 + mScarlet (20 µg)
Pvc13-flg22 (2x GGSGG) + mScarlet (20 µg)
Pvc13-flg22 (4x GGSGG) + mScarlet (20 µg)
Data Points
Best Fit Line
C
Free mScarlet synthesis and purification
Ladder
Flowthrough
Wash
Fold Dilution from concentrate
1x 2x 4x
kDa
250
150
100
75
50
39.8)
37
25
20
15
10
D
free mScarlet
Pvc13-flg22 (ΔPdp1/Pnf)
Brightfield
mScarlet
6 hours
18 hours
30 hours
E
Pvc13-flg22
Pvc13-flg22 (2x GGSGG)
Pvc13-flg22 (4x GGSGG)
Brightfield
mScarlet
6 hours
18 hours
30 hours

**Supplementary Figure 4. Time-course analysis of PVC-mediated mScarlet delivery.** Denaturing Western blot band intensity analyses on **(A)** a serial dilution of the Pvc13-flg22 nanoparticle and **(B)** all PVC variants loaded with HiBiT-tagged mScarlet protein. Band intensities were measured using the FIJI image processing package, and a standard linear regression was performed on a HiBiT standard. **(C)** SDS-PAGE analysis of free NTD-mScarlet protein with bipartite NLS purification via 6xHis-Ni column chromatography. **(D)** Representative confocal images of *Arabidopsis thaliana* protoplasts challenged with 4.5 ng $\mu L^{-1}$ free mScarlet protein and 1 mg $mL^{-1}$ empty Pvc13-flg22 particles. Scale bars, 100 μm. **(E)** Representative confocal images of *Arabidopsis thaliana* protoplasts challenged with 1 mg $mL^{-1}$ mScarlet-loaded Pvc13-flg22, Pvc13-flg22 (2xGGSGG), and Pvc13-flg22 (4xGGSGG) particles. Images were collected at various incubation timepoints. Scale bars, 100 μm.

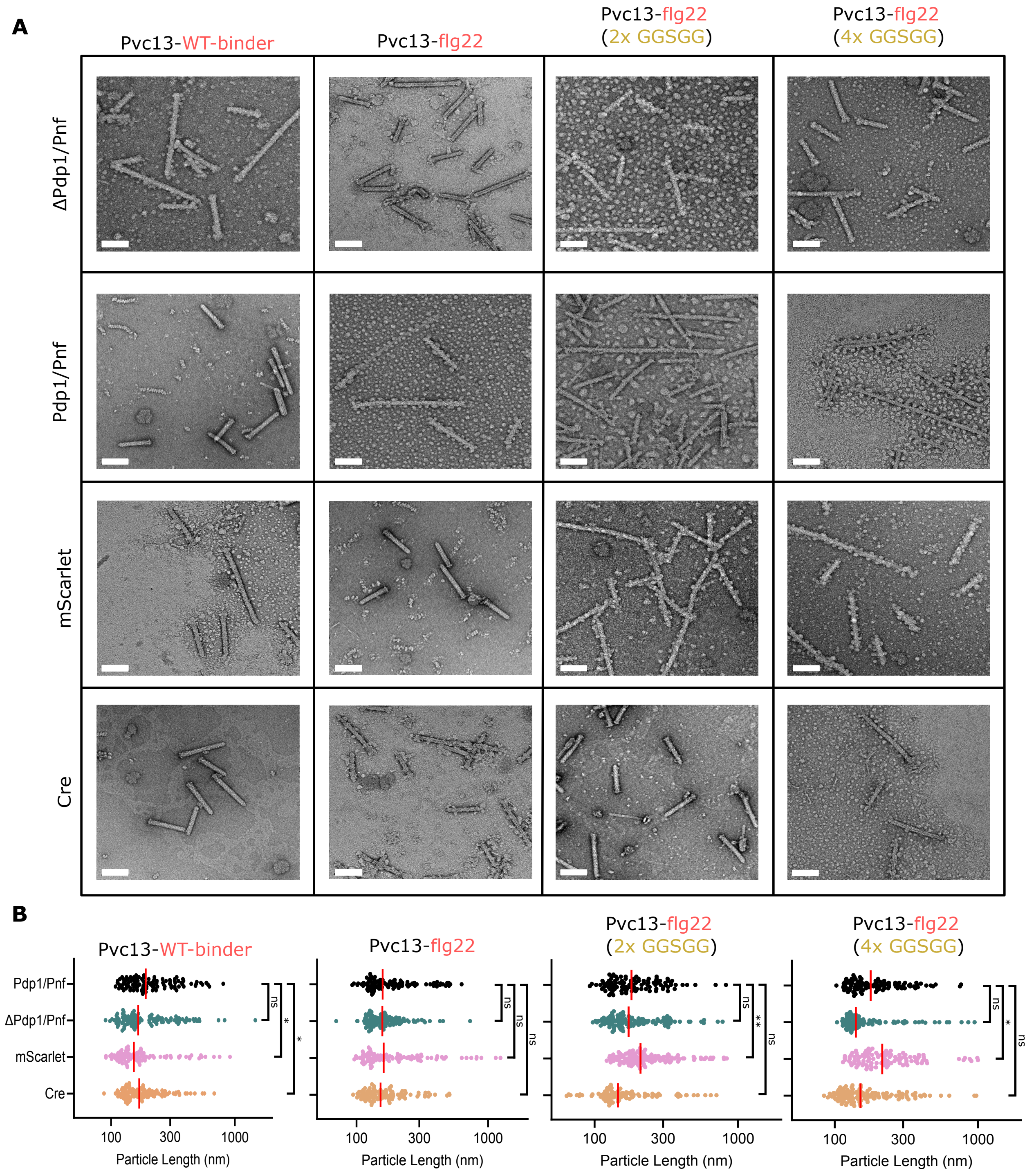


**Supplementary Figure 5. PVCs maintain structural stability upon tail fiber and cargo engineering. (A)** TEM micrographs of all PVC structural and cargo variants. Structural variants are listed in each column, including PVCs with the native WT binding epitope and all three FLS2-binding variants. Cargo variants are listed in each row, including unloaded versions (ΔPdp1/Pnf), native toxin cargos, mScarlet protein, and Cre recombinase. Scale bars, 100 nm. **(B)** Particle length quantification from TEM micrographs, where PVC structural variants are grouped, and their loaded cargoes are permuted. Vertical red bars represent data medians. Data are means with n = 100 individual PVCs counted, Lognormal Brown-Forsythe and Welch's ANOVA with Games-Howell post hoc test. *$p < 0.05$, **$p < 0.01$. ns, not significant.

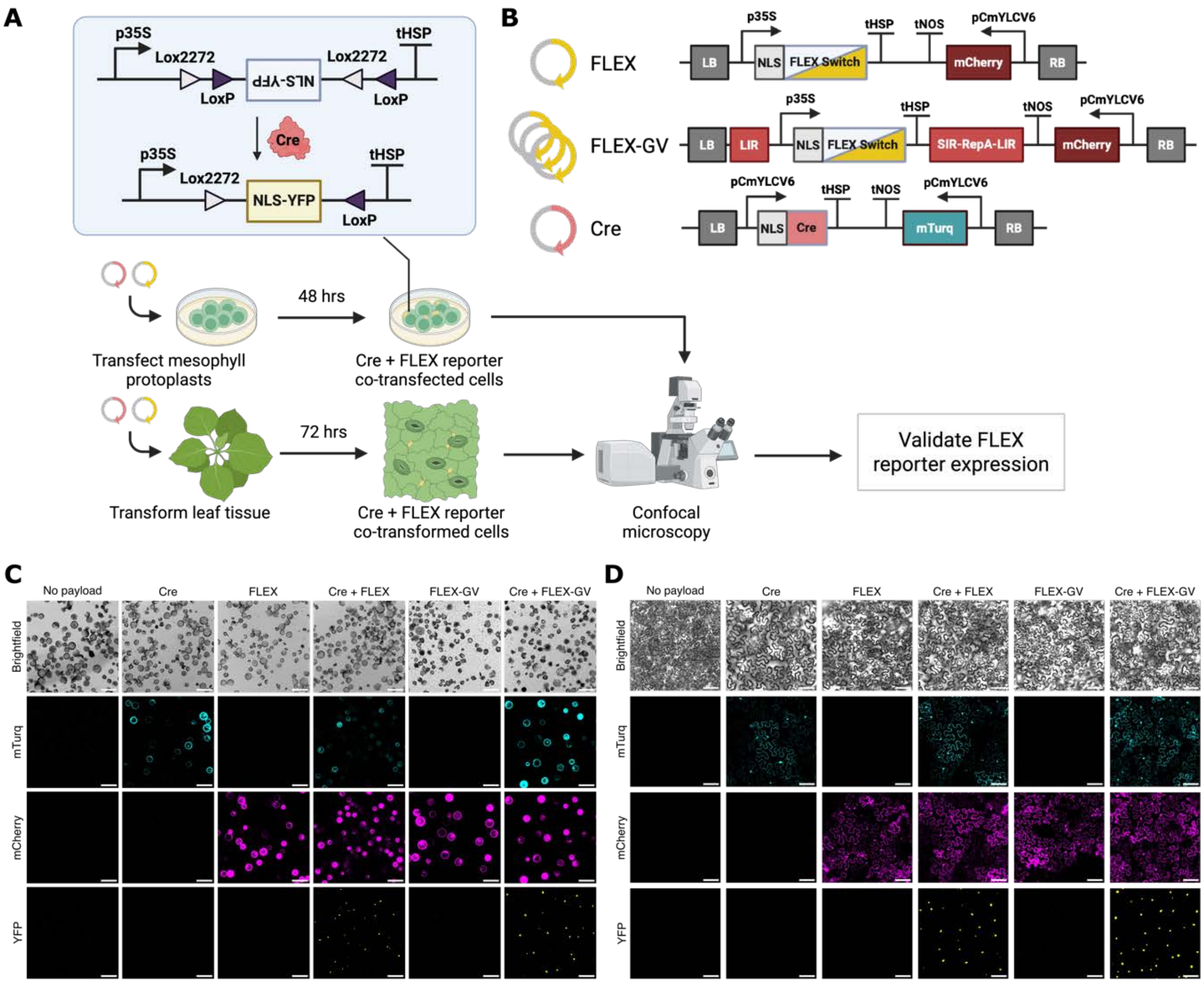


**Supplementary Figure 6. An *in-planta* reporter assay for Cre recombinase delivery detection. (A)** FLEX schematic demonstrating two pairs of orthogonal *loxP*/*lox2272* sites interacting with Cre to achieve irreversible inversion of a reporter sequence. The reporter design produces a nuclear YFP signal in response to Cre activation, enabling visualization of a nuclear-localized signal with confocal microscopy. **(B)** Schematics of two different FLEX reporter designs and a positive control vector. Flanking the minimal FLEX switch with geminiviral components (FLEX-GV) increases reporter copy numbers within harboring cells. A constitutive Cre-expression plasmid serves as a positive control for validating the FLEX switch designs. **(C)** Representative confocal microscopy images of *Arabidopsis thaliana* protoplasts co-transfected with FLEX or FLEX-GV and positive control Cre plasmids. Scale bars, 100 µm. **(D)** Representative confocal microscopy images of *N. benthamiana* leaves co-transformed with FLEX or FLEX-GV and Cre positive control plasmids. Scale bars, 100 µm.

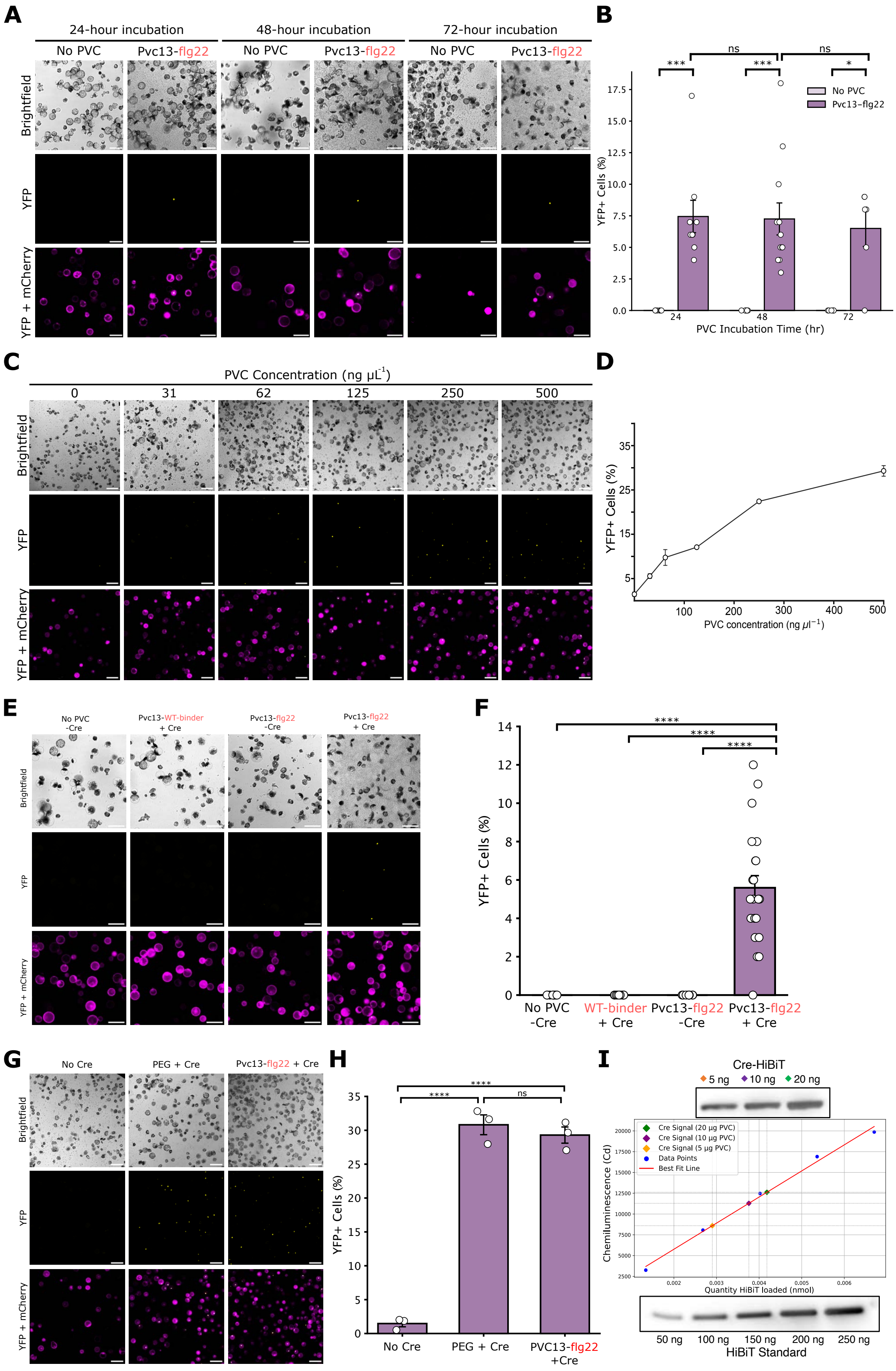

A
24-hour incubation
48-hour incubation
72-hour incubation
No PVC
Pvc13-flg22
Brightfield
YFP
YFP + mCherry
B
***
ns
***
ns
*
No PVC
Pvc13-flg22
YFP+ Cells (%)
0.0
2.5
5.0
7.5
10.0
12.5
15.0
17.5
24
48
72
PVC Incubation Time (hr)
C
PVC Concentration (ng μL-1)
0
31
62
125
250
500
Brightfield
YFP
YFP + mCherry
D
YFP+ Cells (%)
5
15
25
35
100
200
300
400
500
PVC concentration (ng μl−1)
E
No PVC -Cre
Pvc13-WT-binder + Cre
Pvc13-flg22 -Cre
Pvc13-flg22 + Cre
Brightfield
YFP
YFP + mCherry
F
****
****
****
YFP+ Cells (%)
0
2
4
6
8
10
12
14
No PVC -Cre
WT-binder + Cre
Pvc13-flg22 -Cre
Pvc13-flg22 + Cre
G
No Cre
PEG + Cre
Pvc13-flg22 + Cre
Brightfield
YFP
YFP + mCherry
H
****
****
ns
YFP+ Cells (%)
0
5
10
15
20
25
30
35
No Cre
PEG + Cre
PVC13-flg22 +Cre
I
Cre-HiBiT
5 ng
10 ng
20 ng
Cre Signal (20 μg PVC)
Cre Signal (10 μg PVC)
Cre Signal (5 μg PVC)
Data Points
Best Fit Line
Chemiluminescence (Cd)
2500
5000
7500
10000
12500
15000
17500
20000
0.002
0.003
0.004
0.005
0.006
Quantity HiBiT loaded (nmol)
50 ng
100 ng
150 ng
200 ng
250 ng
HiBiT Standard

**Supplementary Figure 7. PVC-mediated Cre delivery in *A. thaliana* protoplasts. (A)** Representative confocal microscopy images of protoplasts harboring the FLEX reporter and challenged with 500 ng $\mu L^{-1}$ Pvc13-flg22 particles. Images were collected at various incubation time points (1-3 days). Scale bars, 100 µm. **(B)** Time-course analysis quantifying %YFP+ cells corresponding to panel A, with PVC incubation time defined as the image acquisition time post-PVC challenge. Data are mean ± SEM with n > 5 technical replicates, one-way ANOVA with Tukey post hoc test. **(C)** Representative confocal microscopy images of protoplasts harboring the FLEX-GV reporter and challenged with various concentrations (0-500 ng $\mu L^{-1}$) of Pvc13-flg22 particles. Images were collected at the 48-h delivery saturation time point. Scale bars, 100 µm. **(D)** Dose-response curve quantifying %YFP+ cells corresponding to panel C. Data are mean ± SEM with n = 3 biological replicates. **(E)** Representative confocal microscopy images of protoplasts harboring the FLEX reporter and challenged with 500 ng $\mu L^{-1}$ of indicated PVC variants. Scale bars, 100 µm. **(F)** Quantification of %YFP+ cells from confocal images in panel F. Data are mean ± SEM with n > 8 technical replicates, one-way ANOVA with Tukey post hoc test. **(G)** Representative confocal microscopy images of protoplasts harboring the FLEX-GV reporter and challenged with Cre protein via PEG transfection or Pvc13-flg22. Scale bars, 100 µm. **(H)** Quantification of %YFP+ cells from confocal images in panel G. Data are mean ± SEM with n = 3 biological replicates, one-way ANOVA with Tukey post hoc test. **(I)** Denaturing Western blot band intensity analysis on a serial dilution of the Pvc13-flg22 nanoparticle loaded with HiBiT-tagged Cre protein. *p < 0.05, ***p < 0.001, ****p < 0.0001. ns, not significant.

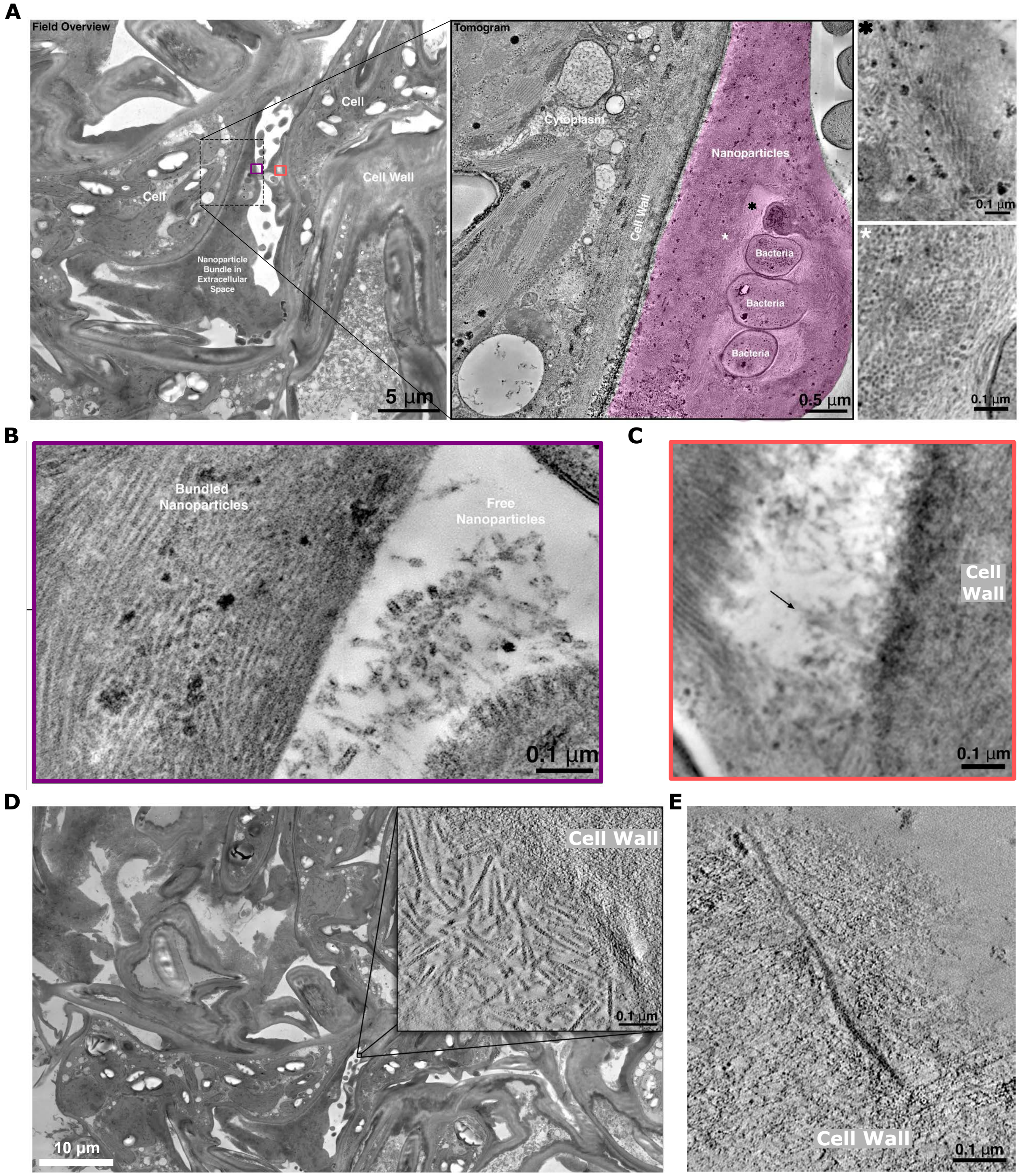


**Supplementary Figure 8. High-resolution TEM images of PVCs in *Nicotiana benthamiana* leaf tissue.** **(A)** TEM field overview of the *N. benthamiana* epidermis 24 hours following Pvc13-flg22 application. Scale bar, 5 µm. The supplemented tomogram demonstrates a region of densely packed PVC nanoparticles proximal to the cell well. Scale bar, 500 nm. *(black) inset visualizes PVC bundles in the longitudinal orientation. *(white) inset visualizes PVC bundles in the cross-sectional orientation. Scale bars, 100 nm. **(B)** A high-resolution image of the purple region in panel A shows individual PVCs adjacent to a dense PVC bundle. Scale bar, 100 nm. **(C)** A high-resolution image of the pink region in panel A shows where PVC bundles are less compact. The black arrow indicates a PVC oriented perpendicular to the cell wall. Scale bar, 100 nm. **(D)** An ultra-high-resolution tomogram directly visualizes individual PVCs within high-density bundles as they interact with the cell wall. Scale bars, 10 µm (100 nm inset). **(E)** A fully extended PVC can be visualized diffusing through the cell wall at regions of lower contrast. Scale bar, 100 nm.

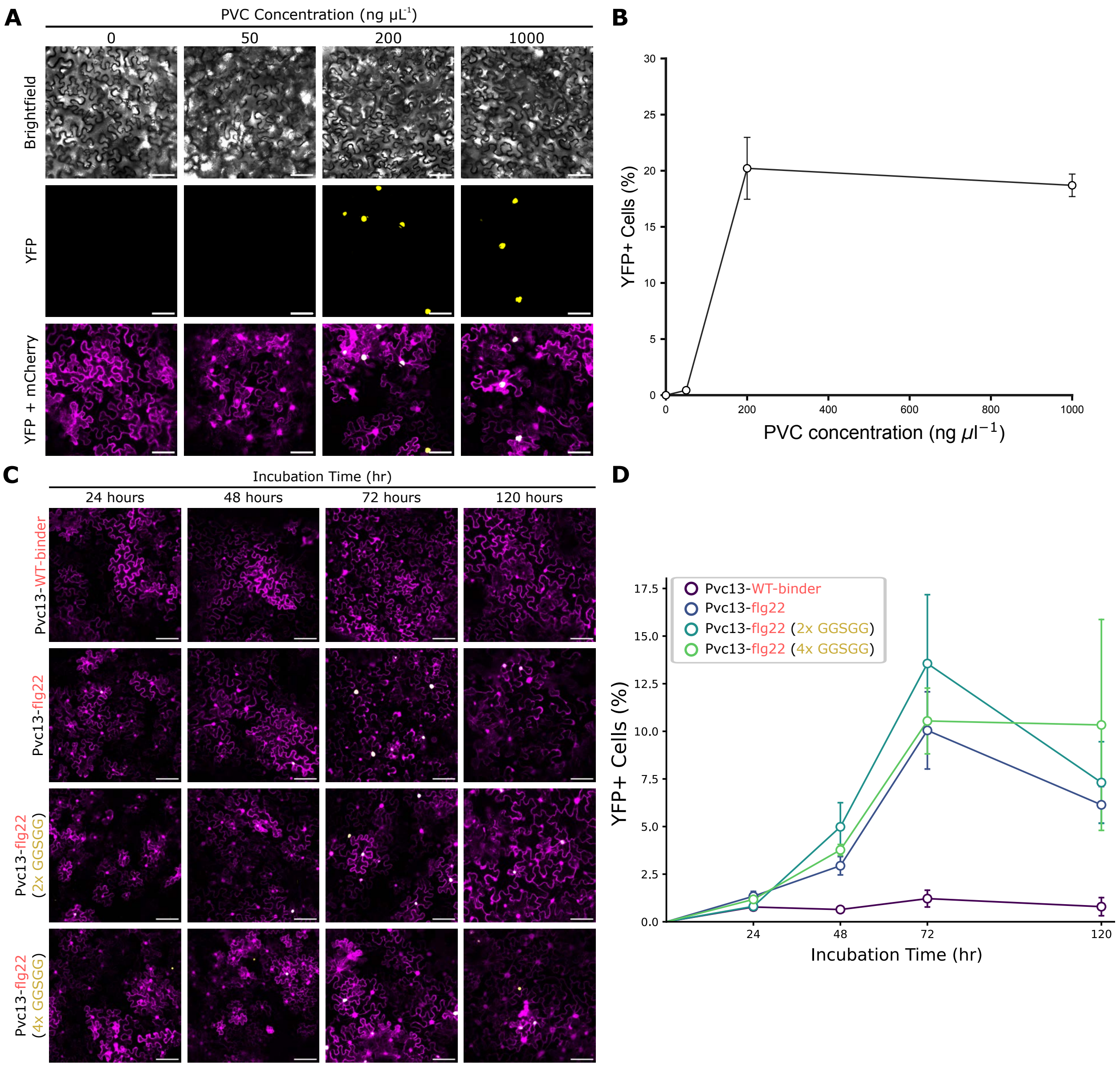


**Supplementary Figure 9. PVC-mediated Cre delivery in *N. benthamiana* leaves. (A)** Representative confocal microscopy images of epidermal leaf tissue harboring the FLEX-GV reporter and challenged with various concentrations (0-1000 ng $\mu L^{-1}$) of Pvc13-flg22 (2xGGSGG). Images were collected at the 72-h delivery saturation time point. Scale bars, 100 µm. **(B)** Dose-response curve quantifying the percentage of YFP+ cells corresponding to panel A. **(C)** Representative confocal microscopy images of epidermal leaf tissue harboring the FLEX-GV reporter and challenged with 500 ng $\mu L^{-1}$ of all PVC variants. Images were collected at various incubation time points (1-5 days). Scale bars, 100 µm. **(D)** Time-course curves quantifying the percentage of YFP+ cells corresponding to panel C, with incubation time defined as the image acquisition time post-PVC challenge. Data are mean ± SEM with n = 4 biological replicates.

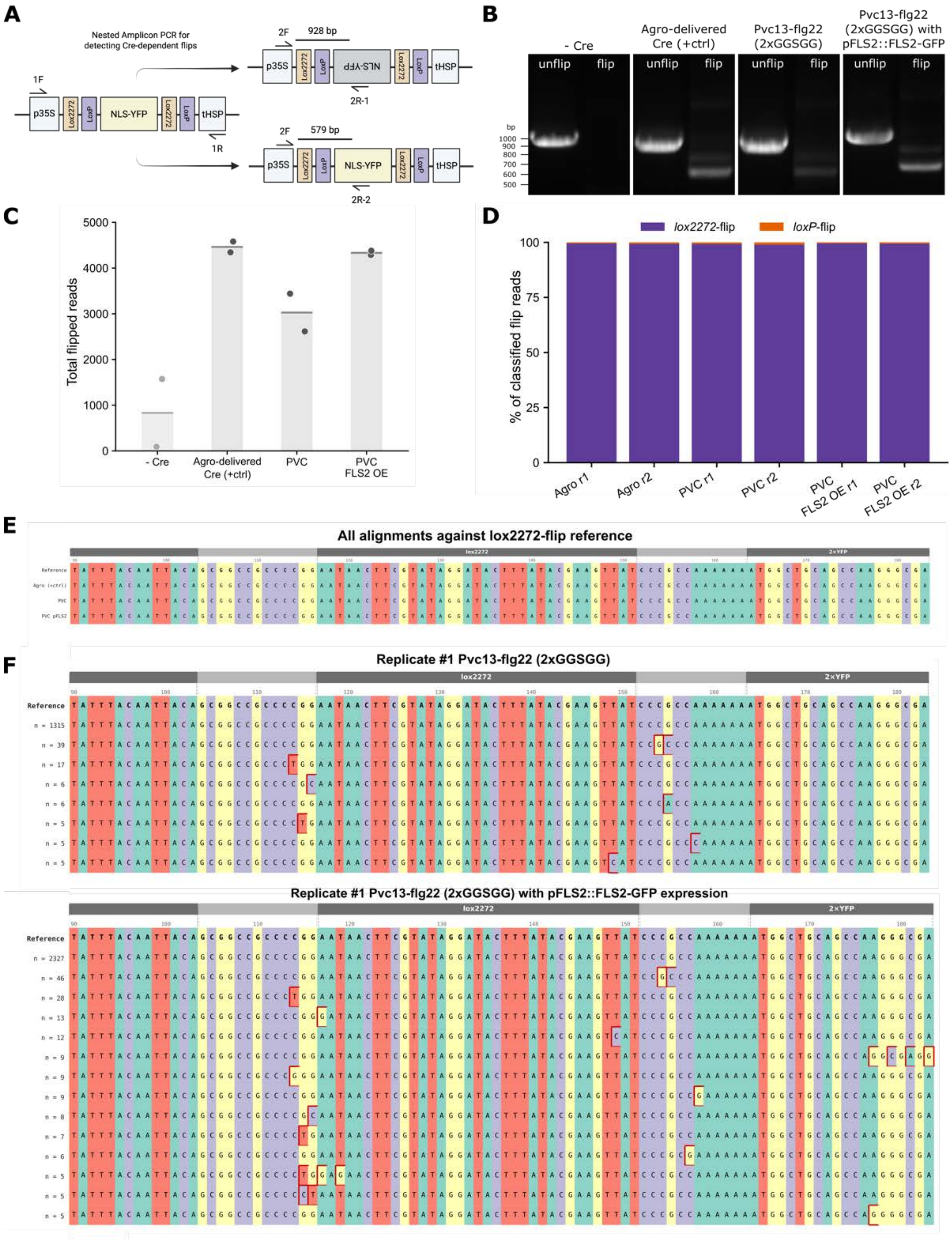

A
Nested Amplicon PCR for detecting Cre-dependent flips
1F
p35S
Lox2272
LoxP
NLS-YFP
tHSP
1R
2F
928 bp
2R-1
579 bp
2R-2
B
- Cre
Agro-delivered Cre (+ctrl)
Pvc13-flg22 (2xGGSGG)
Pvc13-flg22 (2xGGSGG) with pFLS2::FLS2-GFP
unflip
flip
bp
1000
900
800
700
600
500
C
Total flipped reads
5000
4000
3000
2000
1000
0
- Cre
Agro-delivered Cre (+ctrl)
PVC
PVC FLS2 OE
D
lox2272-flip
loxP-flip
% of classified flip reads
100
75
50
25
0
Agro r1
Agro r2
PVC r1
PVC r2
PVC FLS2 OE r1
PVC FLS2 OE r2
E
All alignments against lox2272-flip reference
lox2272
2×YFP
F
Replicate #1 Pvc13-flg22 (2xGGSGG)
Reference
n = 1315
n = 39
n = 17
n = 6
n = 6
n = 5
n = 5
n = 5
Replicate #1 Pvc13-flg22 (2xGGSGG) with pFLS2::FLS2-GFP expression
Reference
n = 2327
n = 46
n = 28
n = 13
n = 12
n = 9
n = 9
n = 9
n = 8
n = 7
n = 6
n = 5
n = 5
n = 5

**Supplementary Figure 10. Molecular validation of FLEX reporter inversion upon PVC-delivered Cre. (A)** Schematic of nested PCR to amplify the 5' lox junctions of the FLEX or FLEX-GV reporters. The nested PCR step employs 2 reverse primers to isolate either the unflipped (2R-1) or the flipped (2R-2) form. **(B)** Nested PCR amplicons from *N. benthamiana* tissue harboring the FLEX-GV reporter and challenged with Cre-loaded Pvc13-flg22 (2xGGSGG) nanoparticles with and without FLS2 overexpression (OE). Benchmarking was performed against the *Agro*-delivered Cre construct used during FLEX reporter validation. **(C)** Total number of flipped amplicon reads (forward primer: 2F, reverse primer: 2R-2) following NGS analysis on flipped amplicons gel extracted from panel B. PVC indicates Cre-loaded Pvc13-flg22 (2xGGSGG) samples; PVC FLS2 OE indicates Cre-loaded Pvc13-flg22 (2xGGSGG) samples with pFLS2::FLS2-GFP expression. Data are means with n = 2 biological replicates. **(D)** Quantification of read alignments against the two expected FLEX inversion-excision products to determine the favored reference sequence. Samples are two biological replicates (r1 and r2) of the conditions from panel B. **(E)** Sequence alignment of each condition from panel B spanning the *lox2272* junction of the preferred FLEX inversion product. **(F)** Multiple sequence alignments of Pvc13-flg22 (2xGGSGG) and Pvc13-flg22 (2xGGSGG) with FLS2 OE samples against the *lox2272* junction of the preferred FLEX inversion product.

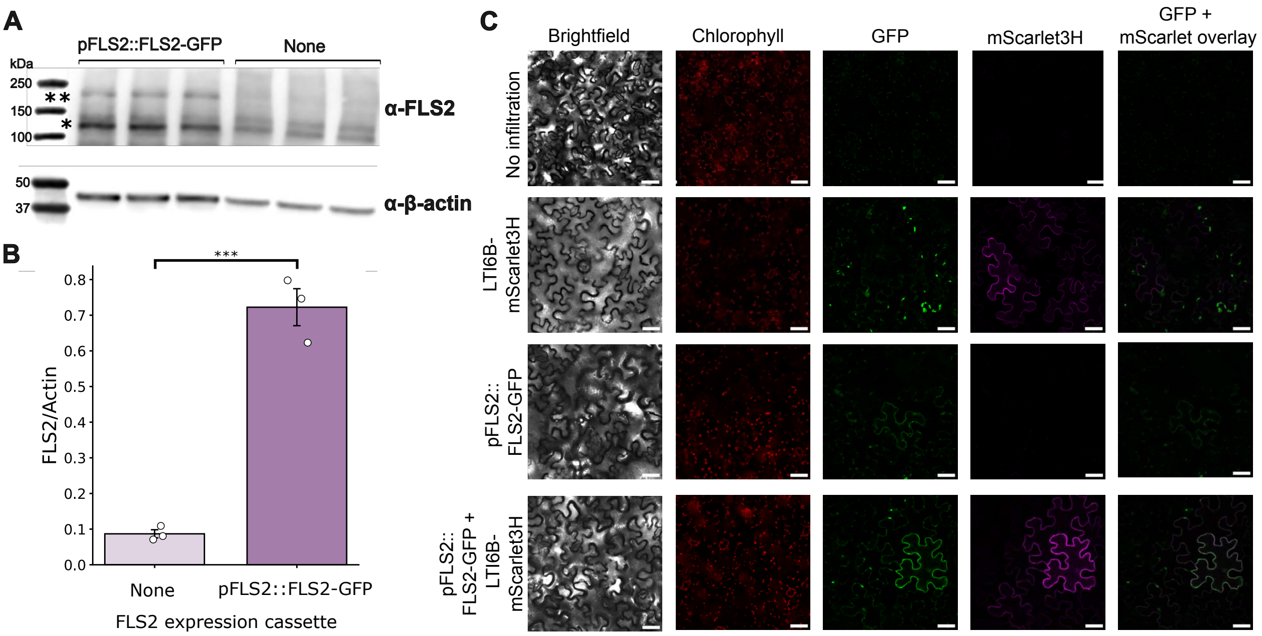


**Supplementary Figure 11. FLS2 overexpression increases receptor abundance at the cell membrane.** **(A)** Western blot of transient pFLS2::FLS2-GFP expression in *N. benthamiana* compared to wild-type leaf homogenates probed with anti-FLS2 antibody. β-actin was used as a loading control for normalization. n = 3 biological replicates per sample. * indicates FLS2, ** indicates overexpressed FLS2-GFP protein. **(B)** Densitometry-based quantification of total anti-FLS2–reactive signal from the blot in panel A, normalized to the corresponding β-actin signal for each lane. Total FLS2 signal was quantified using the gel analysis tool in FIJI by integrating * and ** bands. Data are mean ± SEM with n = 3 biological replicates; one-way ANOVA with Tukey post hoc test. ***$p < 0.001$. **(C)** Representative confocal microscopy images of *N. benthamiana* leaves transiently co-expressing FLS2-GFP constructs and the plasma membrane marker LTI6B-mScarlet3H at 4 days post-*Agrobacterium* infiltration. Chlorophyll autofluorescence was assayed in a separate channel and was distinguishable from the membrane-associated GFP signal. Scale bars, 50 µm.

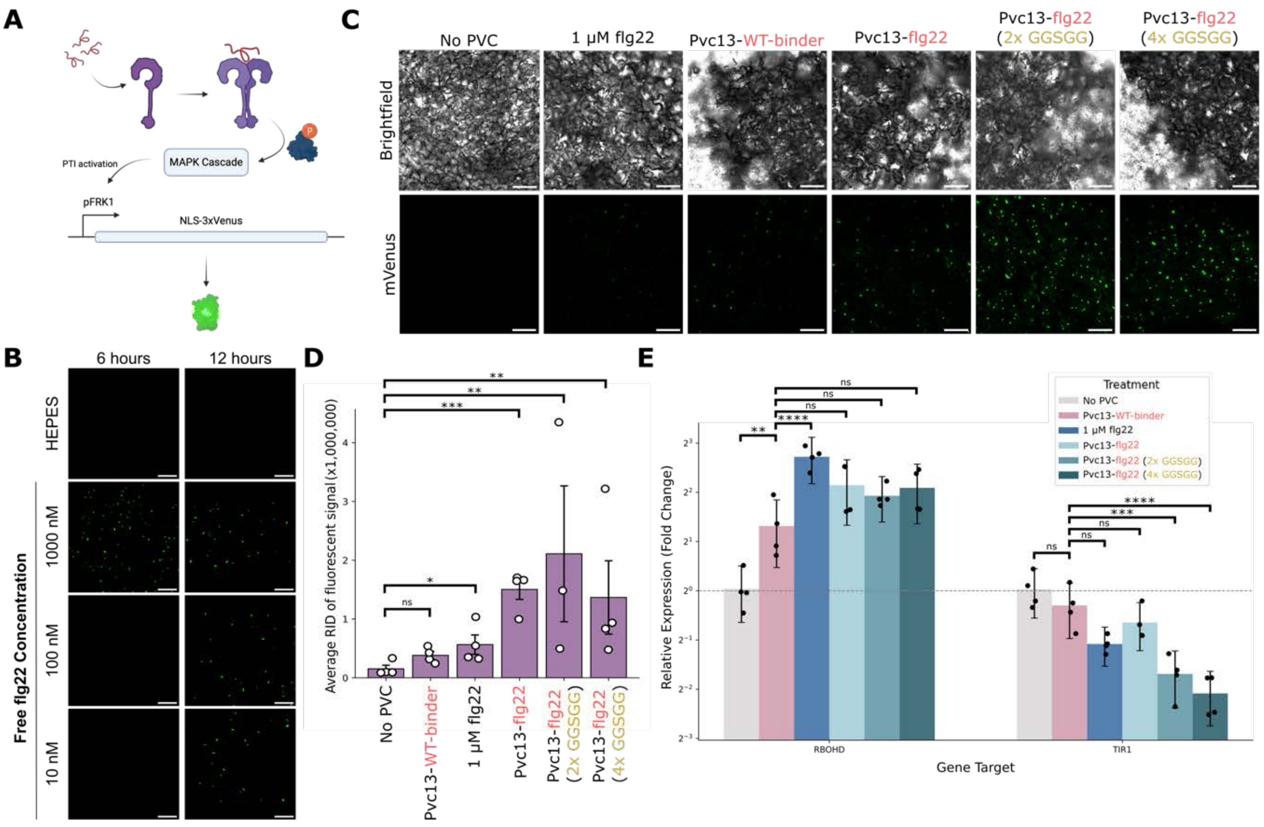


**Supplementary Figure 12. Downstream signaling upon PVC-flg22–FLS2 binding. (A)** Schematic of the downstream MAPK cascade produced upon flg22 binding to the FLS2 receptor and its coreceptor, BAK1, to induce a nuclear mVenus fluorescent reporter in a transgenic pFRK1 *Arabidopsis thaliana* line. **(B)** Representative confocal images of mVenus reporter induction in response to titrations of a synthetic flg22 positive control peptide. Scale bars, 100 µm. **(C)** Representative confocal images of mVenus reporter induction in response to PVC challenge, including a positive free flg22 control and a WT insect-binding control. Scale bars, 100 µm. **(D)** Quantification of total mVenus fluorescence from images collected in panel C, where average RID is the mean raw intensity density across all images of a given biological replicate. **(E)** RT-qPCR analysis of two gene targets, *RBOHD* and *TIR1*, affected by the PVC-flg22–FLS2 binding interaction. Data in panel D are mean ± SEM with n > 3 biological replicates; one-way ANOVA (log-transformed) with Tukey post hoc test. Data in panel E are mean ± SE with n = 4 biological replicates, one-way ANOVA with Tukey post hoc test. Errors were propagated from raw Cq values and transformed into the fold-change space. *p = 0.0672, **p < 0.01, ***p < 0.001, ****p < 0.0001. ns, not significant.

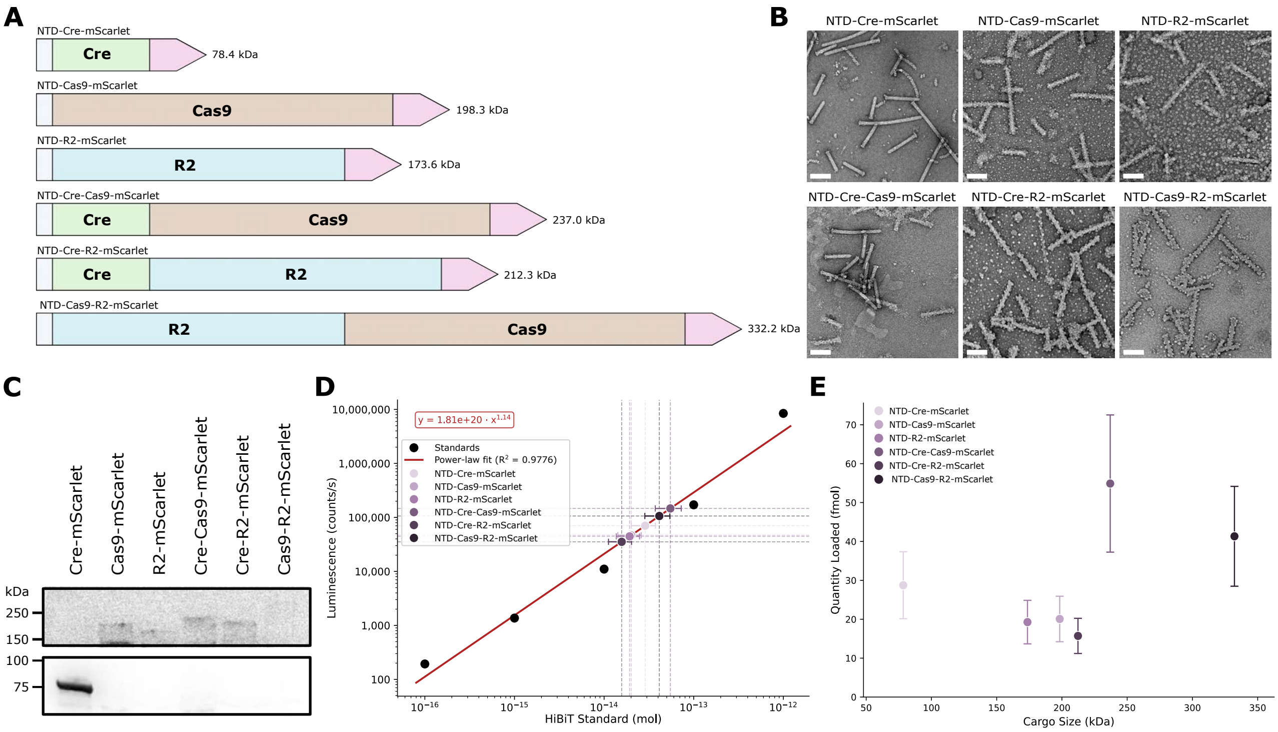


**Supplementary Figure 13. PVCs can load high-molecular-weight protein cargoes. (A)** Cargo fusion protein library of different molecular weights from 78 to 332 kDa. Each protein contains an N-terminal loading tag and a C-terminal mScarlet-HiBiT construct to enable loading detection. **(B)** TEM micrographs of Pvc13-flg22 nanoparticles loaded with each cargo in the variable MW library. Scale bars, 100 nm. **(C)** HiBiT Western blot of the variable cargo library with each PVC formulation loaded at 50 µg except for the NTD-Cre-mScarlet, which was loaded at 10 µg. Transfer of high-MW proteins is more challenging, but faint bands can still be observed on the membrane at increased integration times. **(D)** *In vitro* HiBiT Lytic Assay was performed on denatured PVC formulations from panel B. Variable cargo samples were loaded at a total of 10 µg PVC and quantified against a standard ladder. **(E)** Analysis of cargo loading efficiency from panel D against the total molecular weight of the cargo protein. Data in D and E are from a single replicate, with error propagated from the standard-curve linear fit.
